# Asynchronous Rate Chaos in Spiking Neuronal Circuits

**DOI:** 10.1101/013375

**Authors:** Omri Harish, David Hansel

## Abstract

The brain exhibits temporally complex patterns of activity with features similar to those of chaotic systems. Theoretical studies over the last twenty years have described various computational advantages for such regimes in neuronal systems. Nevertheless, it still remains unclear whether chaos requires specific cellular properties or network architectures, or whether it is a generic property of neuronal circuits. We investigate the dynamics of networks of excitatory-inhibitory (EI) spiking neurons with random sparse connectivity operating in the regime of balance of excitation and inhibition. Combining Dynamical Mean-Field Theory with numerical simulations, we show that chaotic, asynchronous firing rate fluctuations emerge generically for sufficiently strong synapses. Two different mechanisms can lead to these chaotic fluctuations. One mechanism relies on slow I-I inhibition which gives rise to slow subthreshold voltage and rate fluctuations. The decorrelation time of these fluctuations is proportional to the time constant of the inhibition. The second mechanism relies on the recurrent E-I-E feedback loop. It requires slow excitation but the inhibition can be fast. In the corresponding dynamical regime all neurons exhibit rate fluctuations on the time scale of the excitation. Another feature of this regime is that the population-averaged firing rate is substantially smaller in the excitatory population than in the inhibitory population. This is not necessarily the case in the I-I mechanism. Finally, we discuss the neurophysiological and computational significance of our results.

**Author Summary:** Cortical circuits exhibit complex temporal patterns of spiking and are exquisitely sensitive to small perturbations in their ongoing activity. These features are all suggestive of an underlying chaotic dynamics. Theoretical works have indicated that a rich dynamical reservoir can endow neuronal circuits with remarkable computational capabilities. Nevertheless, the mechanisms underlying chaos in circuits of spiking neurons remain unknown. We combine analytical calculations and numerical simulations to investigate this fundamental issue. Our key result is that chaotic firing rate fluctuations on the time scales of the synaptic dynamics emerge generically from the network collective dynamics. Our results pave the way in the study of the physiological mechanisms and computational significance of chaotic states in neuronal networks.

## Introduction

Single cell recordings [1] and electro-encephalography [2, 3] suggest the existence of chaotic dynamics in the brain. Consistent with chaotic dynamics, *in-vivo* experiments have demonstrated that cortical circuits are sensitive to weak perturbations [4, 5]. Remarkably, the misplacement of even a single spike in a cortical network has a marked effect on the timing of subsequent spikes in the network [6].

Chaotic states in extended dynamical systems can be classified as synchronous or asynchronous, depending on the spatial patterns of the dynamics. In synchronous chaos the temporal fluctuations exhibit spatial correlations. If the temporal fluctuations are spatially incoherent, the chaotic state is classified as asynchronous

EEG measures the activity of a large population of neurons. Therefore, it is probable that chaoticity observed in EEGs reflects synchronous chaos in brain regions of rather large size. Models of local cortical circuits exhibiting synchronous chaos have been studied in [7–12]. A computational advantage of synchronous chaos in the brain is that it enables neuronal populations to respond quickly to changes in their external inputs [7] and facilitates the access of the network to states (e.g. limit cycles or fixed points) that encode different stimuli [3]. A large body of experimental data, however, has reported that cortical neurons exhibit very weak correlations [13, 14] and thus are more compatible with asynchronous than with synchronous chaos. Moreover, recent studies have demonstrated that the richness, the complexity and the high dimension of the dynamics in systems operating in asynchronous chaos endows them with remarkable computational capabilities [15–18]. The present paper focuses on the mechanisms underlying the emergence of asynchronous chaos in local neuronal circuits.

Asynchronous chaos was studied in a seminal work by Sompolinsky, Crisanti and Sommers (SCS) [19], who investigated a large network of *N* neuronal-like units fully connected with random weights drawn from a zero mean Gaussian distribution (called hereafter as the SCS model).The dynamics of the network are those of a “rate” model [20], in which the activity of a unit, *S*(*t*), is characterized by a continuous variable which is a non-linear function, *S* = *Φ*(*h*), of the total input to the unit. In the SCS model the activity variables take values between [-1,1] and the function *Φ*(*h*) is sigmoidal and odd. Using Dynamical Mean-Field Theory (DMFT) SCS showed that if the standard deviation of the weight distribution is sufficiently large, the dynamics bifurcate from fixed point to asynchronous chaos. The SCS model in its original form or in its discrete time version has been used in numerous studies in theoretical and computational neuroscience [15–17, 21–25].

However, the connectivity of the SCS model violates Dale’s Law, whereby in biological networks a given neuron is either excitatory or inhibitory [26]. Also, the equation of the SCS model dynamics are invariant under the transformation *h*→−*h*, a symmetry not fulfilled in more realistic neuronal network models. More importantly, as this is the case frequently for rate models, the physiological meanings of the dynamical “neuronal” variables and of the parameters are not clear in the SCS network. Should these variables and the time constant of their dynamics - which sets the time scale of the chaotic fluctuations - be interpreted as characterizing neurons, or synapses?

In this paper we address the following general and fundamental issues: To what extent are asynchronous chaotic states generic in networks of spiking neurons? How does this depend on single neuron properties? How do excitation and inhibition contribute to the emergence of these states? To what extent these chaotic dynamics share similarities with those exhibited by the SCS model? We first study these questions in one population of inhibitory neurons receiving feedforward excitation. We then address them in networks of two populations, one inhibitory and the other excitatory, connected by a recurrent feedback loop. A major portion of the results presented here constitutes the core of the Ph.D thesis of one of the authors (O.H) [27].

## Results

### One population of inhibitory neurons: General theory

We consider *N* randomly connected inhibitory spiking neurons receiving an homogeneous and constant input, *I*. The voltage of each neuron has nonlinear dynamics, as e.g. in the leaky integrate-and fire (LIF model, see *Materials and Methods*) or in conductance-based models [20].

The connection between two neurons is *J*_*ij*_ = *JC*_*ij*_ (*i, j* = 1, 2…*N*), with *J* ≤ 0, and *C*_*ij*_ = 1 with probability *K/N* and 0 otherwise. The outgoing synapses of neuron *j* obey

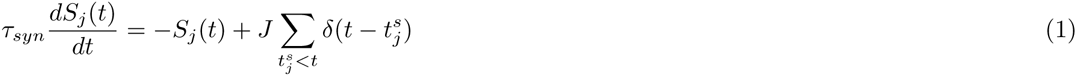

where *S*_*j*_(*t*) is the synaptic current at time *t* and *τ_syn_* the synaptic time constant. When neuron *j* fires a spike 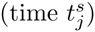, *S*_*j*_ increments by *J*. Thus, the total input to neuron *i*, *h*_*i*_(*t*) = *I* + ∑_*j*_ *J*_*ij*_*S*_*j*_(*t*), satisfies:

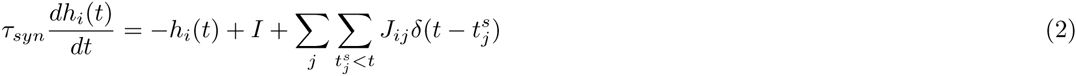

We assume *K* ≫ 1, hence the number of recurrent inputs per neuron is 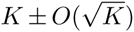. Scaling *J* and *I* as: 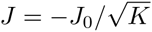, 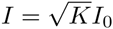, the time-averaged synaptic inputs are 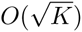 and their spatial (quenched) and temporal fluctuations are *O*(1) [28, 29]. Finite neuronal activity requires that excitation and inhibition cancel to the leading order in *K*. In this *balanced* state, the mean and the fluctuations of the *net* inputs are *O*(1) [28, 29]. The properties of the balanced state are well understood if the synapses are much faster than all the typical time constants of the intrinsic neuronal dynamics [30]. Temporally irregular *asynchronous* firing of spikes is a hallmark of this regime [13,28,29,31,32]. However, this stochasticity does not always correspond to a true chaotic state [28, 29, 33–36]. In fact, this depends on the spike initiation dynamics of the neurons [37]. The opposite situation, in which some of the synapses are slower than the single neuron dynamics, remains poorly understood. This paper mostly focuses on that situation.

When the synaptic dynamics is sufficiently slow compared to the single neuron dynamics, the network dynamics can be reduced to the set of non-linear first order differential equations:

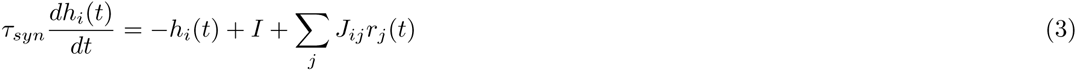

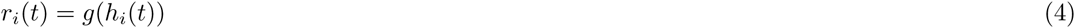

where *r*_*i*_(*t*) is the instantaneous firing rate of neuron *i* and *g*(*h*) is the neuronal input-output transfer function [20]. These are the equations of a *rate model* [20, 39] in which the activity variables correspond to the net synaptic inputs in the neurons. Equations (3)-(4) differ from those of the SCS model in that they have a well defined interpretation in terms of spiking dynamics, the time constant has a well defined physiological meaning, namely, the *synaptic* time constant, the transfer function quantifies the *spiking* response of the neurons and is thus positive, the interactions satisfy Dale’s law and the neuronal connectivity is partial.

#### Dynamical mean-field theory (DMFT)

We build on a DMFT [19] to investigate the dynamics, Eqs. (3)-(4), in the limit 1 ≪ *K* ≪ *N*. Applying this approach, we rewrite the last two terms in the right hand side of Eq. (3) as a Gaussian noise whose statistics need to be self-consistent with the dynamics. This yields a set of self-consistency conditions which determine the statistics of the fluctuations, from which the synaptic net inputs and the firing rates of the neurons can be calculated. This approach is described in detail in the *Materials and Methods* section.

The DMFT shows that, for a given transfer function, depending on the parameters *J*_0_ and *I*_0_, the dynamics either converge to a fixed point state or remain in an asynchronous, time-dependent state. In the fixed point state, the net inputs to the neurons, 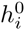, (*i* = 1…*N*) are constant. Their distribution across the population is Gaussian with mean *μ* and variance 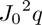. The DMFT yields equations for *μ*, *q*, as well as for the distribution of firing rates 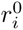 (*i* = 1…*N*) (Eqs. (24)-(25) and (36)). In the time-dependent state, *h*_*i*_(*t*) exhibit Gaussian temporal fluctuations, which are characterized by a mean, *μ* = [⟨*h*(*t*)⟩], and a population-averaged autocovariance (PAC) function, *σ*(*τ*) = [〈*h*(*t*)*h*(*t* + *τ*)〉] − *μ*^2^ ([·] and ⟨.⟩ denote means over the population and over time, respectively). Solving the set of self-consistent equations which determine *σ*(*τ*) and *μ* (Eqs. (25), (27) and (37)-(38)) indicates that *σ*(*τ*) decreases monotonically along the flow of the deterministic dynamics, thus suggesting that the latter are chaotic. To confirm that this is indeed the case one has to calculate the maximum Lyapunov exponent of the dynamics (which characterizes the sensitivity of the dynamics to initial conditions [40]) and verify that it is positive. This can be performed analytically in the framework of DMFT [19]. However, this is beyond the scope of the present paper. Therefore, in the specific examples analyzed below we rely on numerical simulations to verify the chaoticity of the dynamics.

For sufficiently small *J*_0_, the fixed point state is the only solution of the dynamics. When *J*_0_ increases beyond some critical value, *J*_*c*_, the chaotic solution appears. We show in the *Materials and Methods* section that *J*_*c*_ is given by:

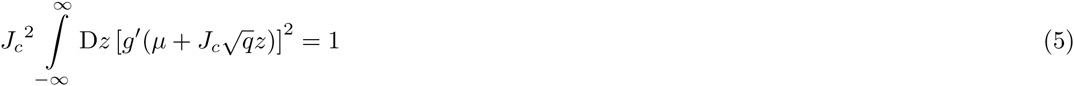

where *q* and *μ* are computed at the fixed point state and 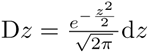.

#### On the stability of the fixed point state

The NxN matrix characterizing the stability of the fixed point is 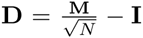 with **I** the NxN identity matrix and:

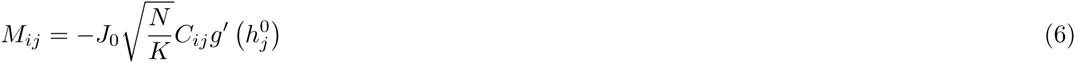

where 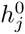 is the total input in neuron *j* at the fixed point. This is a sparse random matrix with, on average, K non zero elements per line or column. In the limit *N ⟶ ∞*, these elements are uncorrelated, have a mean 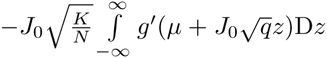 and variance 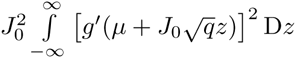(for large *N*, the second moment of the matrix elements is equal to their variance). Interestingly, Eq. (5) means that the SD of the elements of **M** crosses 1 (from below) at *J*_*c*_. As *J*_0_ increases, the fixed point becomes unstable when the real part of one of *v*the eigenvalues crosses 1. Note that that for large *K*, **D** always has a negative eigenvalue, which is 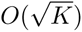.

In the specific examples we investigate below, simulations show that when the chaotic state appears the fixed point becomes unstable. This implies that for *J < J_c_* given by Eq. (5) the real parts of all the eigenvalues of 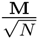 are smaller than 1 and that for *J* = *J*_c_, the real part of one of the eigenvalues, the eigenvalue with maximum real part, crosses 1. This suggests the more general conjecture that in the limit 1 ≪ *K* ≪ *N* the eigenvalue with the largest real part of 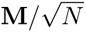 is:

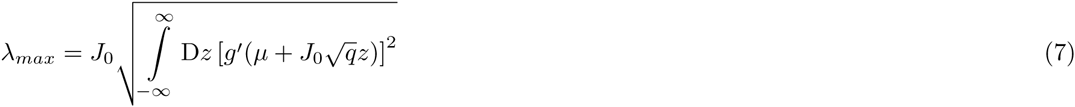

Below we compare this formula to results from numerical diagonalization of 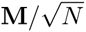.

### One population of inhibitory neurons: Examples

The above considerations show that when synapses are slow, the dynamics of inhibitory networks is completely determined by the transfer function of the neurons. Therefore, to gain insights into the way dynamics become chaotic in such systems we proceed by investigating various spiking models that differ in the shape of their transfer functions.

#### Sigmoidal transfer functions

Neurons in a strong noise environment can be active even if their net inputs are on average far below their noiseless threshold, whereas when these inputs are large the activity saturates. The transfer functions of the neurons can therefore be well approximated by a sigmoid. Specifically here we consider the dynamics described in Eqs. (3)-(4) with a sigmoidal transfer function:

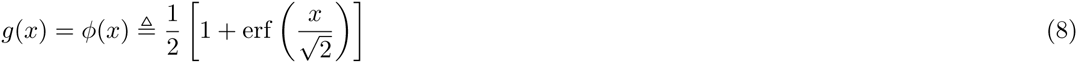

This form of the sigmoid function makes analytical calculations more tractable. Fig. 1A shows that for *J*_0_ =4, *I*_0_=1, the simulated network dynamics converge to a fixed point. This is not the case for *J*_0_ = 6 and *J*_0_ = 15 (Fig 1B,C). In these cases the activities of the neurons keep fluctuating at large time. Note also that the mean level of activity is different for the three neurons. This is a consequence of the heterogeneities in the number of inputs the neurons receive.

**Fig 1.**
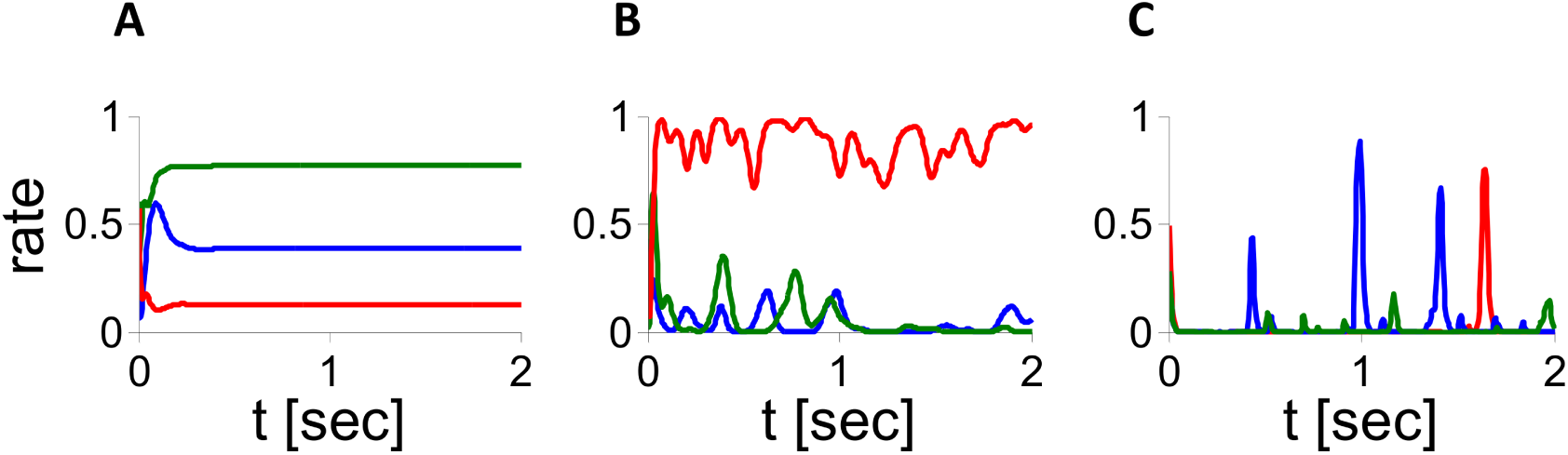
Dynamics in the inhibitory population rate model with *g*(*x*) = *ϕ x*). Activity of 3 neurons in simulations (*N* = 32,000, *K* = 800, *τ_syn_* = 10 ms). A: *J*_0_ = 4. B: *J*_0_ = 6. C: *J*_0_ = 15.

These differences in the network dynamics for these three values of *J*_0_ are consistent with the full phase diagram of the DMFT in the parameter space *I*_0_ − *J*_0_. Fig 2A depicts the results obtained by solving numerically the self-consistent equations that define chaos onset with *g*(*x*) = *Φ* (*x*) (Eqs. (17)-(18) in S2 Text). In the region above the line a chaotic solution exists whereas it does not exist below it. Simulations indicate that in the region above the line, very small perturbations from the fixed point state drive the network toward the time dependent state. In other words, the fixed point solution is unstable above the line: the bifurcation to the time dependent state is thus supercritical.

**Fig 2.**
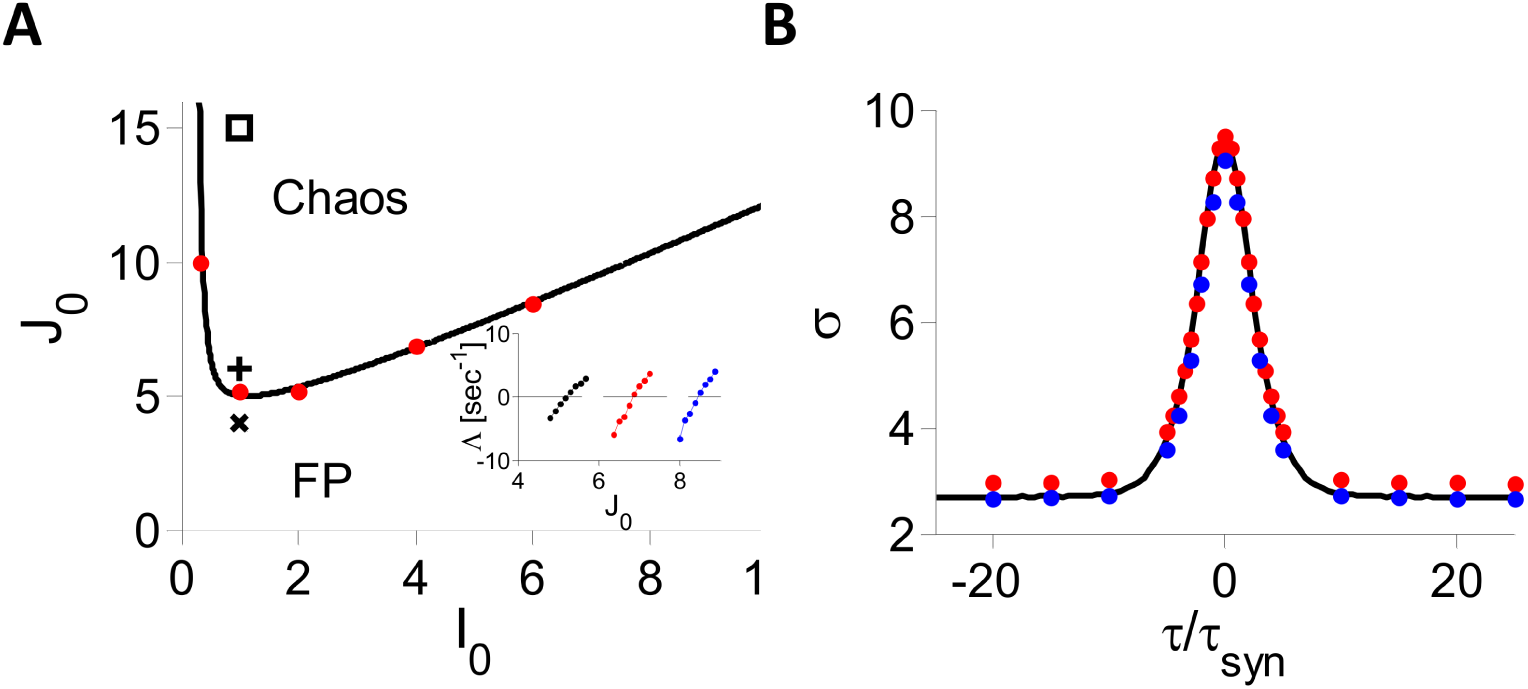
Dynamics in the inhibitory population rate model with *g*(*x*) = *ϕ* (*x*). A: Phase diagram. Solid line: DMFT; Dots indicate where the largest Lyapunov exponent, Λ, changes sign in simulations (*N* = 32,000, *K* = 800, *τ_syn_* = 10 ms). Inset: Λ vs. *J*_0_. *I*_0_ = 2 (black), 4 (red), 6 (blue). Parameters used in Fig.1A,B abd C are marked by ×, + and □, respectively. B: *σ*(*τ*) for *I*_0_ = 1, *J*_0_ = 15. Black: DMFT. Red and blue dots: Simulations for *N* = 32,000, *K* = 800, and *N* = 256,000, *K* = 2000, respectively (results averaged over 8 network realizations).

The instability of the fixed point on this line is also confirmed by direct diagonalization of the matrix 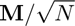 (see Eq. (6)). To this end, we solved numerically the mean field equations for different values of *J*_0_ to obtain *μ* and *q*, randomly sampled 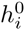 values from the distribution defined by *μ* and *q* to generate the random matrix matrix 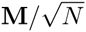, and then computed numerically the spectrum of the matrix (for *N* = 10000). Examples of the results are plotted in Fig 3A for two values of *J*_0_, one below and one above the critical value *J*_*c*_. In both cases, the bulk of the spectrum is homogeneously distributed in the disk of radius *λ*_*max*_ centered at the origin. Fig. 3B plots *λ*_*max*_ computed numerically (dots) and compare the results to our conjecture, Eq. (7) (solid line). The agreement is excellent. The instability of the fixed point corresponds to *λ*_*max*_ crossing 1.

**Fig 3.**
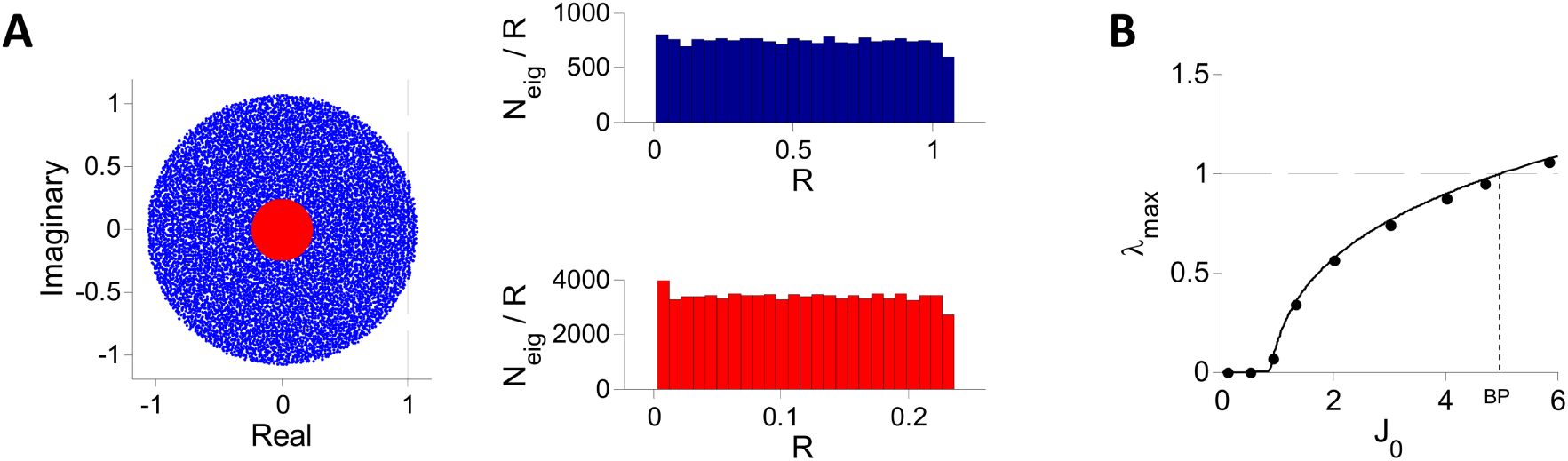
Spectrum of the matrix 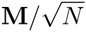 for inhibitory population rate model with (x) = *ϕ* (x). The matrix was diagonalized numerically for *N* = 10000*, K* = 400*, I*_0_ = 1 and different values of *J*_0_. A: The bulk of the spectrum for *J*_0_=6 (blue) and for *J*_0_=1.12 (red). Left: The imaginary parts of the eigenvalues are plotted vs. their real parts for one realization of **M**. This indicates that the support of the spectrum is a disk of radius *λ*_*max*_. Right: Histograms of *N*_*eig*_/*R* (one realization of **M**) where *N*_*eig*_ is the number of eigenvalues with a modulus between *R* and *R* + Δ *R* (Δ *R* = 0.0428 (top), 0.0093 (bottom)) for *J*_0_ = 6 (top) and *J*_0_ = 1.12 (bottom). The distribution of eigenvalues is uniform throughout the spectrum support. B: The largest real part of the eigenvalues (black dots), *λ_max_*, is compared with the conjecture, Eq. (7) (solid line). The fixed point loses stability when *λ_max_* crosses 1.

To verify the chaoticity of the time dependent state predicted by the DMFT in the region above the bifurcation line we simulated the dynamics and computed numerically the largest Lyapunov exponent, Λ, for different values of *I*_0_ and *J*_0_ (see *Materials and Methods* for details). The results plotted in Fig 2A (red dots and inset) show that Λ crosses zero near the DMFT bifurcation line and is positive above it. Therefore the dynamics observed in simulations are chaotic in the parameter region above this line as predicted by the DMFT.

We solved numerically the parametric self-consistent differential equation which determined the PAC, *σ*(*τ*), (Eqs. (25), (29) and (37)-(38)) for different values of *J*_0_ and *I*_0_. An example of the results is plotted in Fig 2B. It shows that numerical simulations and DMFT predictions are in very good agreement. Moreover, simulations with increasing values of *N* and *K* indicate that the small deviations from the DMFT predictions are due to finite *N* and *K* effects; a detailed study of these effects is reported in S1 Text.

Fig. 4A shows the bifurcation diagram of the PAC amplitude, *σ*_0_ − *σ*_∞_. For *J*_0_ below the bifurcation point (BP) the PAC amplitude is zero, which corresponds to the fixed point state (solid blue line). At the bifurcation the fixed point loses stability (dashed blue line) and a chaotic state with a strictly positive PAC amplitude emerges (black line).

**Fig 4.**
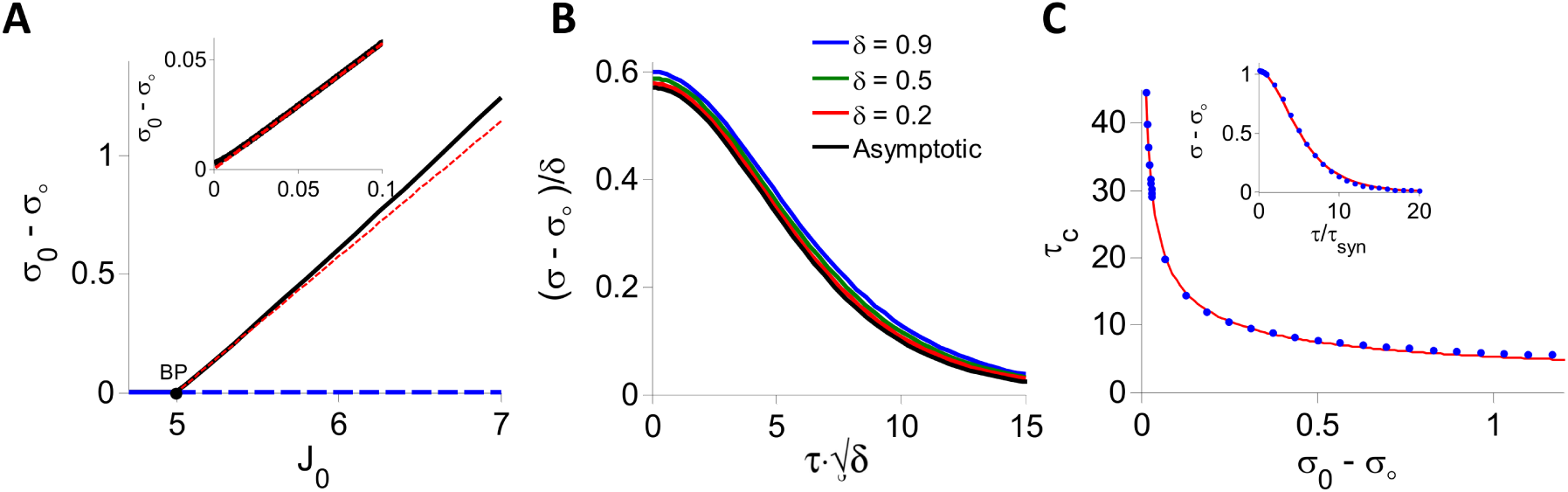
DMFT for the inhibitory rate model with *g*(*x*) = *ϕ* (*x*), *I*_0_ = 1. A: The PAC amplitude, *σ*_0_ − *σ*_*∞*_, is plotted against *J*_0_. At fixed point *σ*_0_ − *σ*_*∞*_ = 0 (blue). When *J*_0_ = *J*_*c*_ *≈* 4.995 (black dot, BP) the chaotic state appears. For *J*_0_ > *J*_*c*_, the fixed point is unstable (dashed blue) and the network settles in the chaotic state (*σ*_0_ *- σ _∞_ >* 0, black). Red: Perturbative solution in the limit *J*_0_ *→ J_c_* (see S2 Text). Inset: *σ*_0_ − *σ*_*∞*_ vanishes linearly when *δ* = *J*_0_ *- J_c_ →* 0 ^+^. Black: Numerical solution of the DMFT equations. Red: Perturbative solution at the leading order, *O*(*δ*). B: (*σ σ ^∞^*)*/ δ* is plotted for different values of *δ >* 0 showing the convergence to the asymptotic form (Eq. (11) in S2 Text) in the limit *d →* 0. C: Blue dots: Decorrelation time, *τ_dec_* vs. PAC amplitude. The PAC, *σ*(*τ*) − *σ*_*∞*_, was obtained by solving numerically the DMFT equations and *τ_dec_* was estimated by fitting the result to the function *A/cosh* ^2^(*τ /τ_dec_*). Red In the whole range, *J*_0_ *∊* [5, 7] considered, *τ_dec_* can be well approximated by 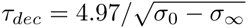. This relation becomes exact in the limit *σ*_0_ *- σ_∞_ →* 0. Inset: Numerical solution of the DMFT equations for *J*_0_ = 6.65 (blue dots) and the fit to *A/cosh* ^2^(*τ/τ_dec_*) (red). The fit is very good although this is far from bifurcation.

We studied analytically the critical behavior of the dynamics at the onset of chaos. We solved perturbatively the DMFT equations for 0 < *δ* = *J*_0_ *- J_c_ ≪* 1, as outlined in the *Materials and Methods* section and in S2 Text. This yields (*σ*(*τ*)-*σ _∞_*) ∝ *δ ^α^*/cosh^2^ (*τ/τ_dec_*), with *α* = 1 and a decorrelation time scaling like *τ*_dec_ ∝ *δ ^β^* with *β* = *-*1*/*2. Therefore at the onset of chaos, the PAC amplitude vanishes and the decorrelation time diverges. We show in the *Materials and Methods* section that this critical behavior with exponents *α* = 1, *β* = −1*/*2, is in fact a general property of the model, Eqs. (3)-(4), whenever *g*(*h*) is twice differentiable. It should be noted that in the SCS model the PAC also vanishes linearly at chaos onset. However, the critical exponent of the decorrelation time is different (*β* =−1) [19].

The inset in Fig 4A compares the PAC amplitude obtained by numerically solving Eq. (27) (black line) with the corresponding perturbative result (red line) for small *δ*. The agreement is excellent. In fact, the perturbative calculation provides a good estimate of the PAC even if *δ* is as large as 0.2*J*_*c*_ (Fig 4A, main panel and Fig 4B). More generally, the PAC can be well fitted with the function (*σ*_0_ -σ_*∞*_)·cosh^-2^ (*τ* /*τ_dec_*) (Fig.4C, inset) providing an estimate of the decorrelation time, *τ_dec_*, for all values of *J*_0_. Fig. 4C plots *τ_dec_* vs. *σ*_0_ *- σ _∞_* for *I*_0_ = 1. It shows that the formula 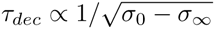 we derived perturbatively for small *δ* provides a good approximation of the relationship between the PAC amplitude and the decorrelation time even far above the bifurcation.

#### Threshold power-law transfer function

We next consider the dynamics of the network (Eqs. (3)-(4)) with a transfer function

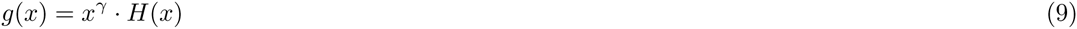

where γ > 0 and *H*(*x*) = 1 for *x >* 0 and 0 otherwise. Non-leaky integrate-and-fire neurons [41] (see also S3 Text) and *θ*-neurons [42–45] correspond to γ = 1 and *γ* = 1*/*2, respectively. The transfer functions of cortical neurons *in-vivo* can be well fitted by a power-law transfer function with an exponent *γ* ≈ 2 [46,47].

Fig. 5A plots the phase diagrams in the *J*_0_ - *I*_0_ parameter space by solving the DMFT equations (see S4 Text) for different values of γ > 1*/*2. For fixed *I*_0_, *J*_*c*_ varies non-monotonically as *γ* decreases. This non-monotonicity is also clear in Fig 5B. When γ ⟶ (1*/*2)^+^, *J*_*c*_⟶ 0 as 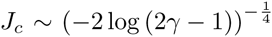 as we show analytically in S4 Text. For γ < 1*/*2, the integral in the right hand side of Eq. (5) diverges. Equivalently, the elements of the stability matrix have infinite variance. Therefore, the DMFT predicts a chaotic dynamics as soon as *J*_0_ *>* 0.

**Fig 5.**
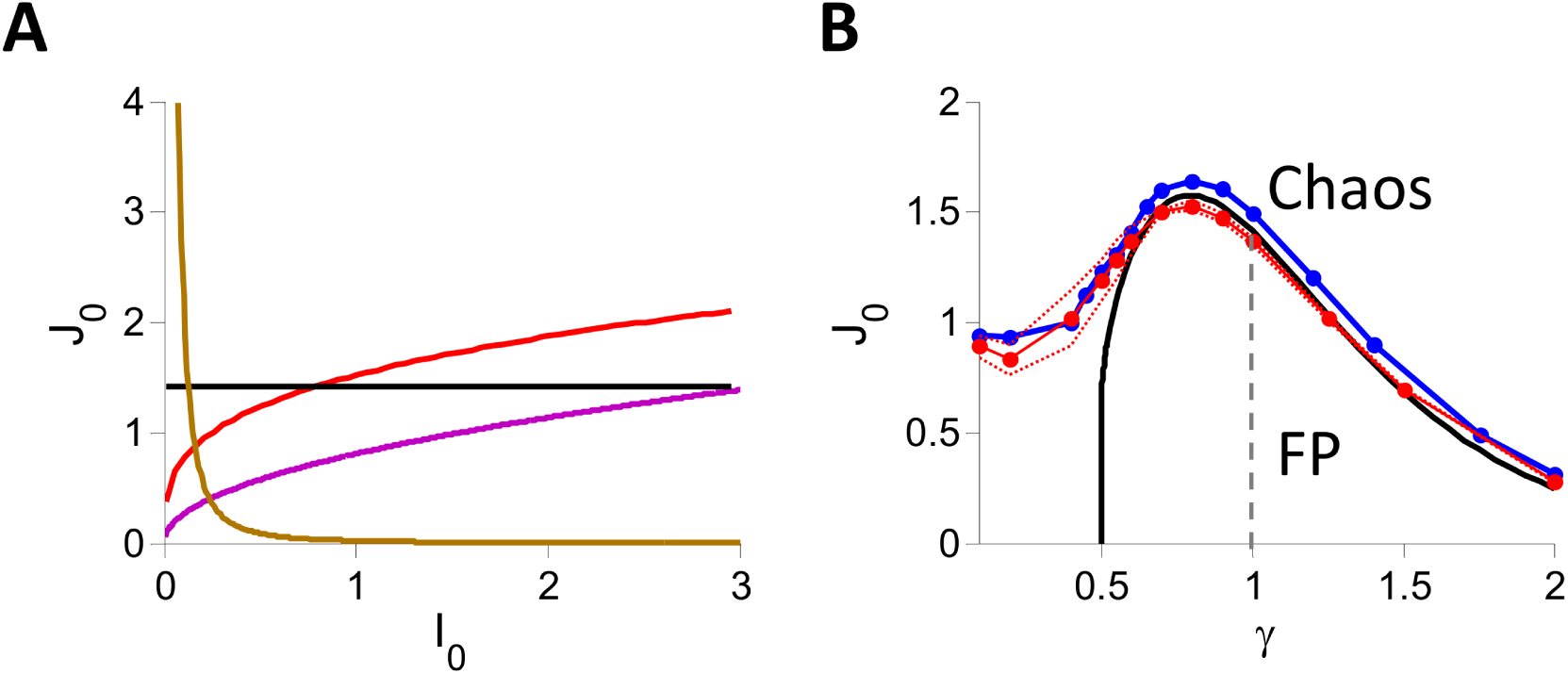
Phase diagrams of inhibitory rate models with *g*(*x*) = *x ^γ^ H*(*x*), *K* = 400. A: *γ* = 3 (gold), 1 (black), 0.7 (red), 0.51 (purple). B: *J*_*c*_ vs. *γ* for *I*_0_ = 1. Black: DMFT. Blue and red: Simulations with *N* = 32000, *K* = 400. Blue: Zero-crossing of λ. Red: The fraction of networks with stable fixed point is 50%, 5% and 95% on the solid, bottom-dashed and top-dashed lines respectively.

To compare these predictions with numerical simulations, we simulated different realizations of the network (N=32000, K=400, *I*_0_ = 1) for various values of *J*_0_. For each value of *J*_0_ and γ we determined whether the dynamics converge to a fixed point or to a time dependent state as explained in the *Materials and Methods* section. This allowed us to compute the fraction of networks for which the dynamics converge to a fixed point. The solid red line plotted in Fig 5B corresponds to a fraction of 50% whereas the dotted red lines correspond to fractions of 5% (upper line) and 95% (lower line). We also estimated the Lyapunov exponent, Λ, for each values of *J*_0_ and γ. The blue line in Fig 5B corresponds to the location where Λ changes sign according to our estimates (see *Materials and Methods* for details).

For 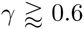, the fraction of networks with an unstable fixed point varies sharply from 0 to 100% in the vicinity of the bifurcation line predicted by the DMFT. Moreover, for these values of γ, the spectrum of the matrix 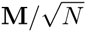 is homogeneously distributed in the disk of radius *λ_max_* centered at the origin and the values of *λ_max_* agrees with Eq. (7). This is shown in Fig 6A for *γ* = 1. Finally, simulations indicate that the values of *J*_0_ where the largest Lyapunov Λ becomes positive in numerical simulations (blue line in Fig 5B) are very close to the DMFT bifurcation values.

However, as γ →(1*/*2)^+^, the discrepancies between DMFT and simulations become more pronounced. Very close to γ = (1*/*2)^+^ there is a whole rage of values of *J*_0_ for which the DMFT predicts chaos whereas in numerical simulations the dynamics always converge to a fixed point. This discrepancy can be understood by observing that the integral over the Gaussian measure in Eq. (5) corresponds to a population average over neurons. When γ → (1*/*2)^+^, the region where *z* is just above -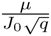 dominates the integral; in other words, the neurons with positive close-to-threshold net inputs are those that make the largest contribution to the destabilization of the fixed point. On the other hand, the DMFT shows that these neurons become extremely rare as γ → (1*/*2)^+^: in that limit *μ_c_* increases sharply, thus shifting the center of the Gaussian distribution to very large positive values. Therefore, we would need to simulate outrageously large networks to obtain a quantitative agreement with the DMFT predictions for the locations of the bifurcation to chaos. Similar arguments explain why when γ < 1*/*2 we find a transition from fixed point to chaos in numerical simulations for 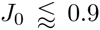 although according to the DMFT the fixed point is always unstable since the integral in Eq. (5) diverges.

**Fig 6.**
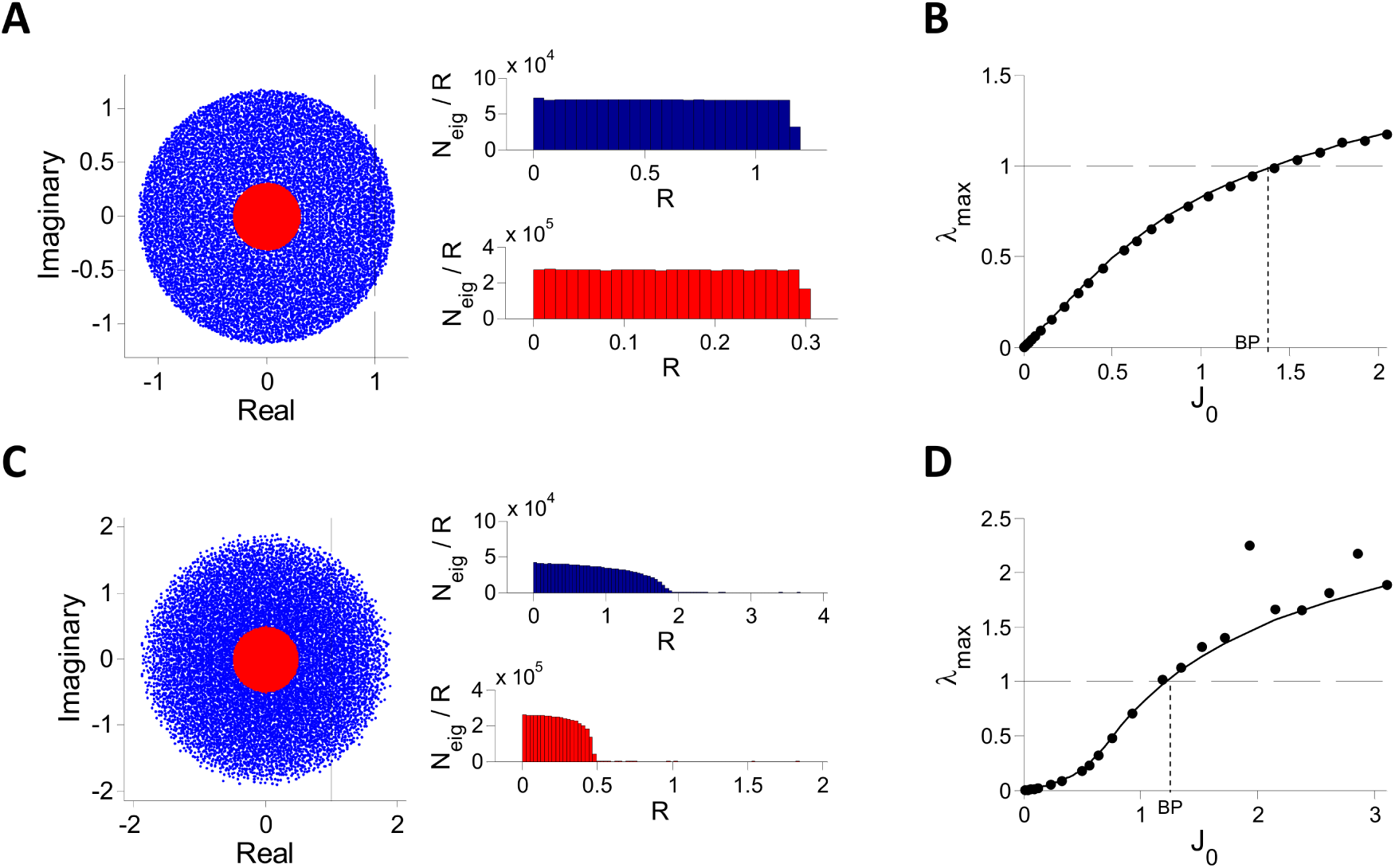
Spectrum of the matrix 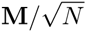 for inhibitory rate models with *g*(*x*) = *x ^γ^ H*(*x*). A-B: *γ* =1. The matrix was diagonalized numerically for *N* = 10000*, K* = 400*, I*_0_ = 1 and different values of *J*_0_. A: The bulk of the spectrum (one realization). Left panel: Imaginary vs. real parts of the eigenvalues for one realization of **M**. Blue: *J*_0_ = 2.045. Red: *J*_0_=0.307. Right panel: Histograms (100 realizations) of *N*_*eig*_/*R* where *N*_*eig*_ is the number of eigenvalues with modulus between *R* and *R* + Δ *R* (Δ *R* = 0.0479 (top), 0.0122 (bottom)) for *J*_0_ =2.045 (top) and *J*_0_=0.307 (bottom). The eigenvalues are almost uniformly distributed throughout the disk of radius *λ_max_* (except very close to the boundary). B: The largest real part of the eigenvalues, *λ_max_* (one realization, black dots) is compared with the conjecture Eq. (7) (solid line). C,D: Same as in A, B, for *γ* = 0.55. Blue: *J*_0_=3.01, Δ *R* = 0.0491; red: *J*_0_ = 0.75, Δ *R* = 0.0246 (red). The agreement with Eq. (7) is good for *J*_0_ not too large but the eigenvalues distribution is non-uniform. Quantitatively similar results are found for *N* = 20000*, K* = 400 as well as *N* = 40000*, K* = 600 (not shown).

Numerical diagonalization of 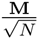 shows that when 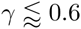 (i) the eigevalues in the bulk of the spectrum are distributed in a disk centered at the origin and that this distribution is less and less homogeneous as γ → (1*/*2)^+^ (ii) the eigenvalue *λ_max_* governing the instability exhibit substantial deviations from Eq. (7) especially for large *J*_0_ (Fig 6C) (iii) *λ_max_* exhibits large sample to sample fluctuations (results not shown). We conjecture that these features are due to large finite *N* and *K* effects and stem from the fact that the SD of the elements of 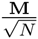 diverges when γ → (1*/*2)^+^.

We studied the dynamics in detail for γ = 1. The DMFT predicts that 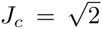 for all *I*_0_ and *K* (*K* large). As already mentioned, the simulations agree well with this result (Fig5B). We studied analytically the dynamics for *J*_0_ close to this transition (Fig.7A-C). To this end, we solved the self-consistent DMFT equations in the limit *δ* = *J*_0_ - *J*_*c*_ → 0^+^. The perturbative calculation, explained in S4 Text, is less straightforward than in the case of a sigmoid transfer function. This stems from the fact that at the threshold, the threshold-linear transfer function is only differentiable once. It yields that *σ*; - σ_∞_ ∼ δ^α^ *σ_s_*(*τ /δ^β^*) with *α* = 2, *β* = -1/2 and the function *σ_s_*(*x*)) has to be determined numerically. The function *σ_s_* is plotted in Fig 7B. It can be well fitted to the function *A* [cosh(*x/x_dec_*)]^-1^ with *A*=12.11 and *x*_*dec*_=2.84 (see Fig 7B, inset). In particular, for small *δ*, the amplitude and the decorrelation time of the PAC are related by *τ_dec_* ∝ 1*/*(*σ*_0_ − *σ _∞_*)^1/4^. Note that the amplitude of the PAC vanishes more rapidly (*α* = 2) than for sigmoidal transfer functions (*α* = 1) whereas the decorrelation time diverges with the same critical exponent (*β* = -1*/*2) in the two cases.

**Fig 7.**
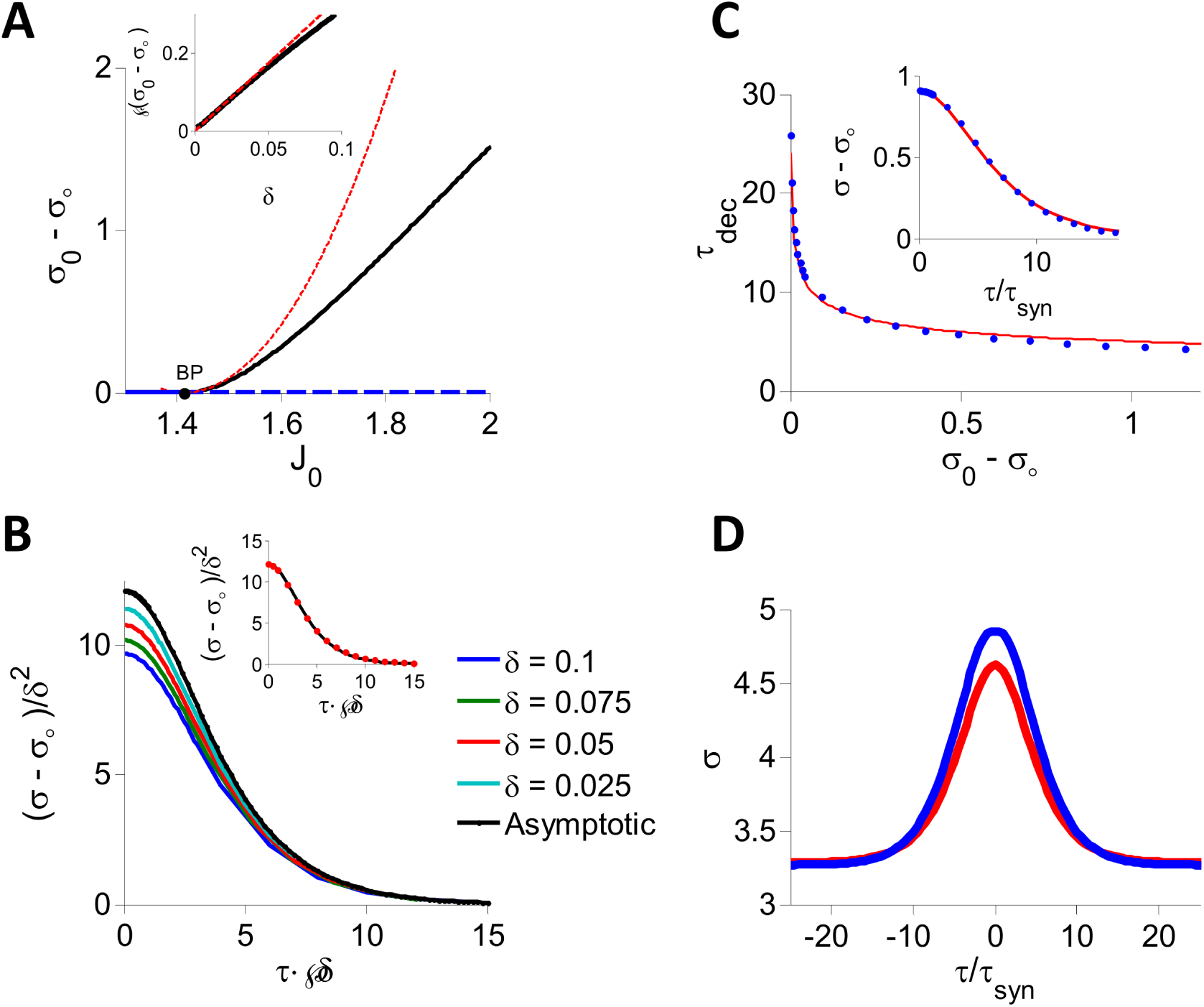
DMFT for the inhibitory rate model with threshold-linear transfer function. A: The PAC amplitude, *σ*_0_ − *σ*_*∞*_, is plotted against *J*_0_. At fixed point *σ*_0_ *- σ_∞_* = 0 (blue). When 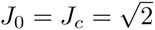 (black dot, BP) a bifurcation occurs and the chaotic state appears. For *J*_0_ *> J_c_*, the fixed point is unstable (dashed blue) and the network settles in the chaotic state (*σ*_0_ *- σ_∞_ >* 0, black). Red: Perturbative solution in the limit *J*_0_ *→ J_c_* (see S4 Text). Inset: 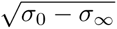 plotted against *δ* = *J*_0_ *- J_c_* showing that *σ*_0_ − *σ*_*∞*_ vanishes quadratically when *d →* 0 ^+^. Black: Full numerical solution of the DMFT equations. Red: Perturbative solution at the leading order, *O*(*δ*). B: (*σ σ_∞_*)*/ δ* ^2^ is plotted for different values of *δ >* 0 to show the convergence to the asymptotic function derived perturbatively in S4 Text. Inset: The function (*σ*(*τ*) - *σ_∞_*)/ *δ* ^2^ (black) can be well fitted to *A/* cosh(*x/x_dec_*) (red dots, *A* = 12.11, *x*_*dec*_ = 2.84). C: Decorrelation time, *τ_dec_* vs. PAC amplitude (blue). The function *σ*(*τ*) - *σ^∞^* was obtained by integrating numerically Eq. (29) and *τ_dec_* was estimated by fitting this function to *A/* cosh(*τ/τ_dec_*). Red: In the whole range of *J*_0_ considered (*J*_0_ ∈ [1.4, 1.9] the relation between *τ_dec_* and *σ*_0_ *- σ_∞_* can be well approximated by 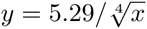. Inset: The PAC computed by solving the DMFT equations for *J*_0_ = 1.81 (blue dots) and the fit to 0.93*/* cosh(*τ /*4.6). D: The PAC for *J*_0_ = 2 and *K* = 1200. Blue: Numerical integration of Eq. (29). Red: Numerical simulations for *N* = 256,000.

Fig. 7A-C compares the results of the perturbative analysis to those of the numerical integration of the differential equation, Eq. (27). Unlike what we found for the sigmoid transfer function, *δ* must be very small 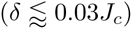 to achieve a good quantitative agreement. It should be noted, however, that the quality of the fit of *σ - σ* _∞_ to *A* [cosh(*τ /τ_dec_*)]^-1^ does not deteriorate by much even far from the bifurcation (Fig 7C, inset; *δ* = 0.4), and that the relation 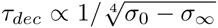 holds with good approximation even if *δ* is not small (Fig 7C, main panel).

Finally, Fig 7D compares DMFT and numerical simulations results for *σ*(*τ*) when *J*_0_ = 2. The agreement is reasonably good but not perfect. We show in S1 Text that the discrepancy between the two improves as the network size increases but that finite size effects are stronger here than in the rate model with sigmoid transfer function.

#### Leaky integrate-and-fire (LIF) inhibitory networks

Our objective here is to obtain further insights into the relevance of the chaotic behavior exhibited by rate dynamics, Eqs. (3)-(4), to understand *spiking* network dynamics. The dynamics of one population of LIF spiking neurons reduces to Eqs. (3)-(4) with the transfer function

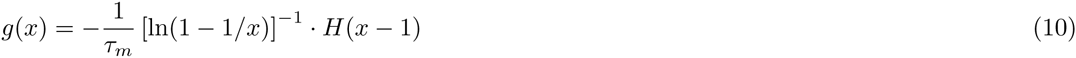

in the limit where the synapses are much slower than the cell membrane time constant, *τ_m_*. Our goal is twofold: 1) to study the emergence of chaos in this rate LIF rate model and 2) to compare it to full spiking dynamics and characterize the range of values of the synaptic time constant for which the two dynamics are quantitatively or at least qualitatively similar.

Fig.s 8,9 depict typical patterns of neuronal activity in simulations of the inhibitory spiking LIF model. For strong and fast synapses (*τ_syn_* =3 ms, Fig. 8A), neurons fire spikes irregularly and asynchronously (Fig.8A). Fig. 8B shows that when *τ_syn_*=100 ms the population average firing rate remains essentially the same (∼ 14.1 Hz) and the network state stays asynchronous. The spiking patterns, however, change dramatically: with large *τ_syn_* neurons fire irregular bursts driven by slowly decorrelating input fluctuations (Fig 9A, blue). Fig. 9B shows that reducing *J*_0_ increases the firing rate, reduces the amplitude of the fluctuations (Fig.9B, inset) and slows down their temporal decorrelation. Eventually, for small enough *J*_0_, *σ*(*τ*) becomes flat and the fluctuations are negligible.

**Fig 8.**
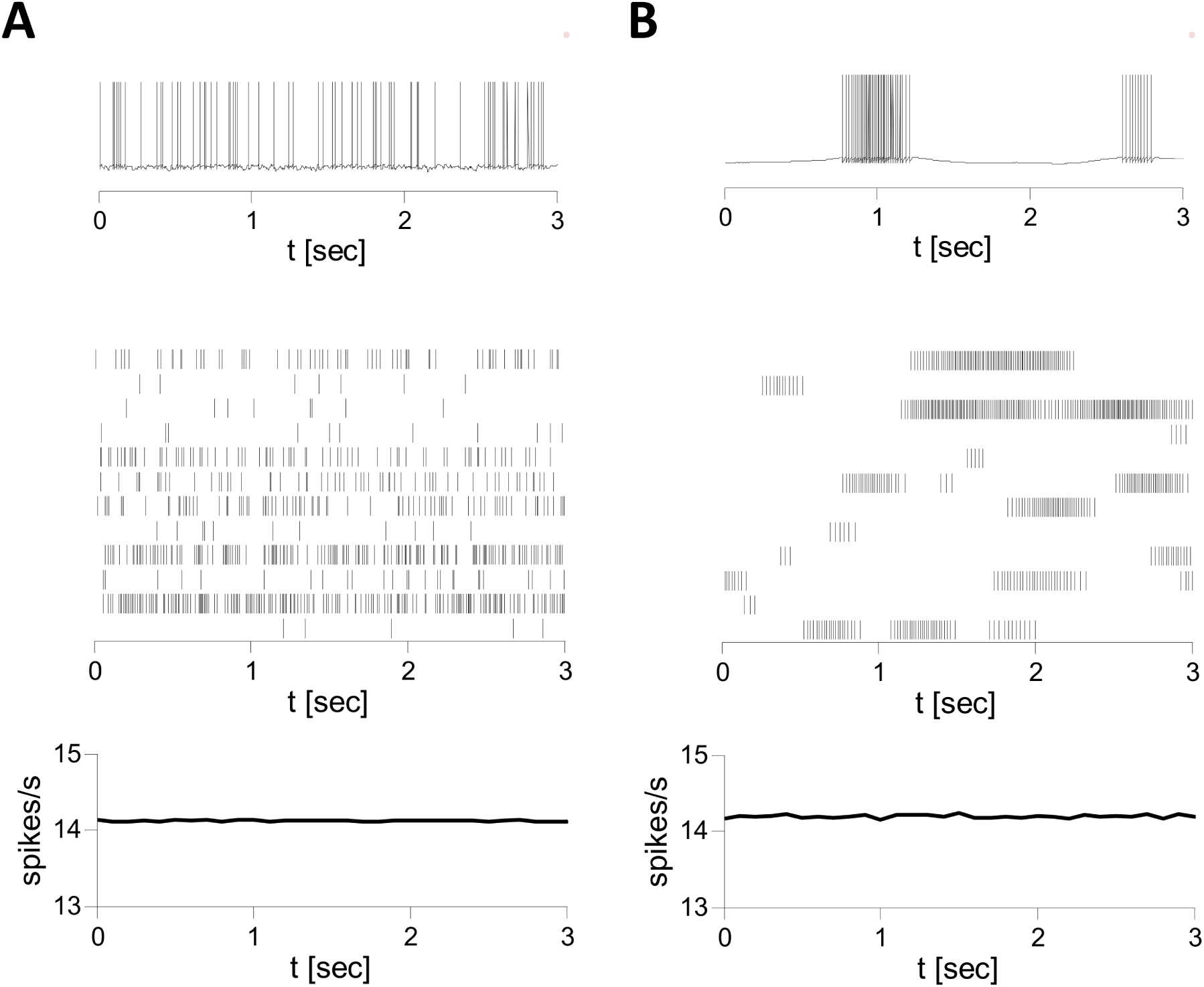
Patterns of activity in simulations of the LIF inhibitory spiking network. *N* = 10000, *K* = 800, *J*_0_ = 2, *I*_0_ = 0.3. Voltage traces of single neurons (top), spike trains of 12 neurons (middle) and population averaged firing rates (in 50 ms time bins, bottom) are plotted. A: *τ_syn_* = 3 ms. Neurons fire irregular spikes asynchronously. B: *τ_syn_* = 100 ms. Neurons fire bursts of action potentials in an irregular and asynchronous manner.

**Fig 9.**
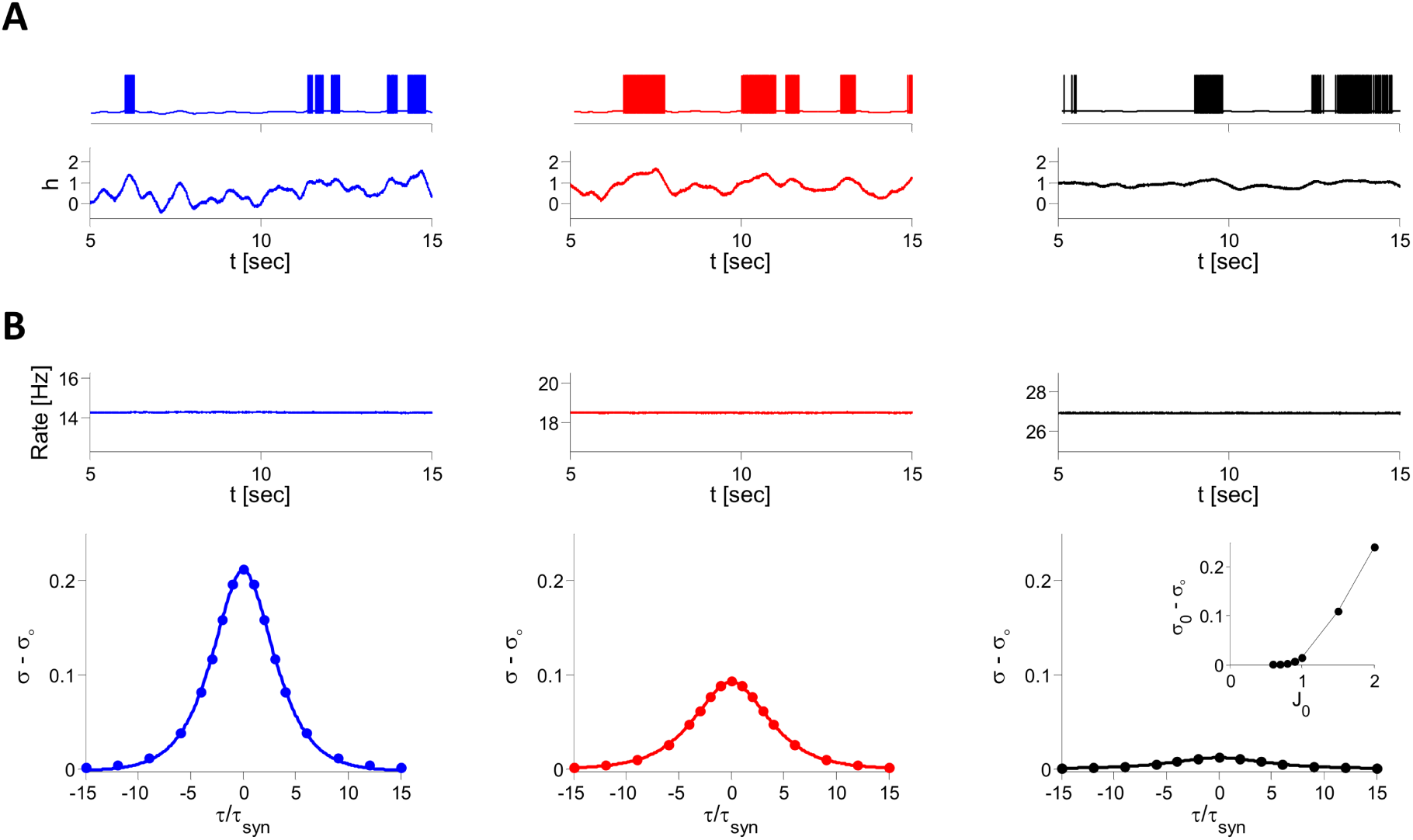
Dependence of the dynamics on synaptic strength in the LIF inhibitory spiking model. Simulation results for *N* = 40,000, *K* = 800, *I*_0_ = 0.3, *τ_syn_* = 100 ms. From left to right: *J*_0_ = 2 (blue), 1.5 (red) and 1 (black). A: Examples of single neuron membrane voltages (top) and net inputs, *h*, (bottom). For the three values of *J*_0_, the mean firing rate of the example neuron is 11 Hz. As *J*_0_decreases, the temporal fluctuations in the net input become smaller whereas the temporal average increases such that the firing rate remains nearly unchanged. B. Top: Population average firing rate increases like 100*I*_0_*/J*_0_ as implied by the balance condition. Bottom: PAC (*σ - σ_∞_* bottom). The dots correspond to the fit of the PAC to (*σ*_0_ − *σ*_*∞*_) *·* [cosh (*τ /τ_dec_*)] ^-1^ which yields *τ_dec_/τ_syn_* = 2.5 (blue), 3.0 (red), 3.8 (black) for the three values of *J*_0_.Inset in the right panel: *σ*_0_ − *σ*_*∞*_ vs. *J*_0_.

**Fig 10.**
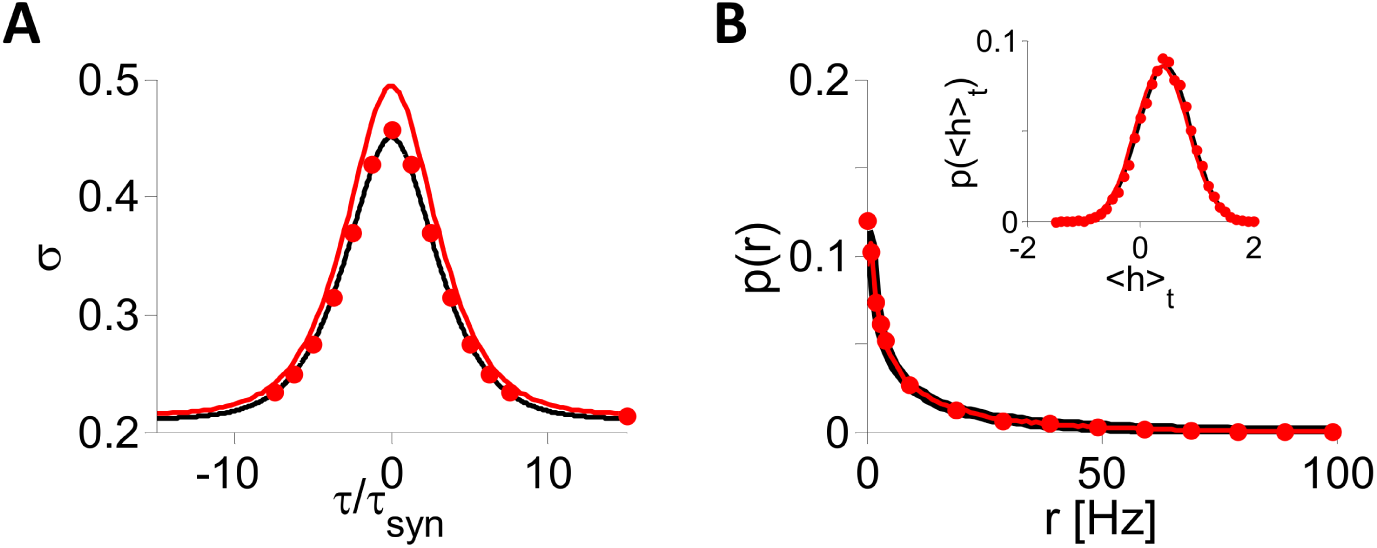
Comparison of the inputs and firing rate statistics in the inhibitory LIF spiking and rate models (simulations and DMFT). *N* = 40,000, *K* = 800. *J*_0_ = 2, *I*_0_ = 0.3, *τ_syn_* = 100 msec. A: *σ*(*τ /τ_syn_*). B: Distributions of neuronal mean firing rates, 〈*r*_*i*_〉, and net inputs, 〈*h*_*i*_〉 (inset) in the spiking network (black) and rate model (red; dots: simulations, solid line: DMFT).

Fig. 10 compares the dynamics of the rate to those of the spiking LIF networks. Panels A,B show that for *J*_0_ = 2*, I*_0_ = 0.3 and *τ_syn_* = 100 ms, *σ*(*τ*), the distributions of the time averages of neuronal firing rates and net inputs, 〈*r*_*i*_〉 and 〈*h*_*i*_〉, are essentially the same in the simulations of the two networks. When reducing *τ_syn_* down to 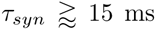, the function *σ*(*τ /τ_syn_*) measured in the spiking network simulations, changes only slightly. In fact, this function is remarkably similar to what is found for the corresponding function in the DMFT and in simulations of the LIF rate model (Fig 11A). Fitting *σ*(*τ*) with the function *B* + *A* [cosh(*τ /τ_dec_*)]^-1^ yields *τ_dec_* ≈ 2.45 *· τ_syn_*.

**Fig 11.**
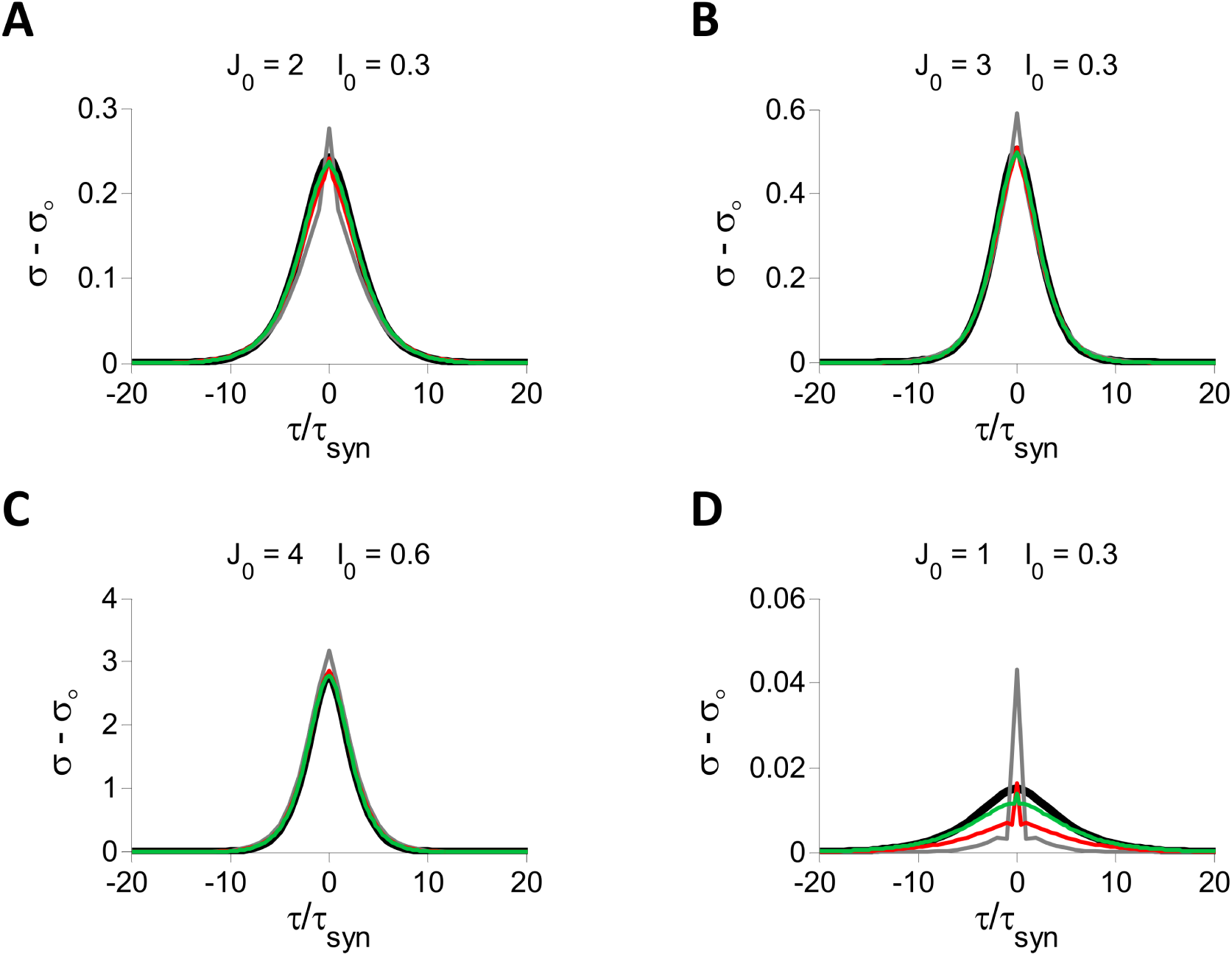
PACs in inhibitory LIF spiking and rate models. All the results are from numerical simulations with *N* = 40,000, *K* = 800. A: *J*_0_ = 2*, I*_0_ = 0.3. B: *J*_0_ = 3*, I*_0_ = 0.3. C: *J*_0_ = 4*, I*_0_ = 0.6. D: *J*_0_ = 1*, I*_0_ = 0.3. In all four panels the PACs are plotted for the spiking network with *τ_syn_* = 10 (gray), 20 (red) and 40 (green) ms. The results for the rate model are also plotted (black). The firing rates are ∼15 Hz in A and C, ∼10 Hz in B and ∼30 Hz in D, in good agreement with the prediction from the *⟨ ⟩* balance condition ([〈 *r 〉*] = 100*I*_0_*/J*_0_ Hz). As the population firing rate increases, a larger *τ_syn_* is needed for good agreement between the spiking and the rate model.

How small can *τ_syn_* be for the two models to still behave in a quantitatively similar manner? Simulations show that this value increases with the mean activity of the network (see examples in Fig 11) but that for reasonable firing rates, fewer than several several tens of Hz, the fluctuations have similar properties in the two models even for *τ_syn_* ≈ 20 ms.

We conducted extensive numerical simulations of the inhibitory LIF rate and spiking models (*N* = 40000, *K* = 800) to compute their phase diagrams in the *I*_0_ - *J*_0_ parameter space. The results for the rate model are plotted in Fig 12. For sufficiently small *J*_0_ the dynamics always converge to a fixed point whereas for sufficiently large *J*_0_ the network always settles in a state in which the activity of the neurons keeps fluctuating at large time. We show in S5 Text that in this regime the maximum Lyapunov exponent is strictly positive, therefore the dynamics are chaotic. Between these two regimes, whether the dynamics converge to a fixed point or to a chaotic state depends on the specific realization of the connectivity matrix. The fraction of networks for which the convergence is to a fixed point depends on *J*_0_. The range of *J*_0_ where this fraction varies from 95% to 5% is notably large as shown in Fig 12. Additional simulation results on this issue are depicted in S5 Text. The counterpart of this behavior in the spiking network is that when *J*_0_ is small, neurons fire regular spikes tonically whereas for sufficiently large *J*_0_ they fire highly irregular bursts. The transition between the two regimes occurs for similar values of *J*_0_ in the rate and in the spiking networks. In both networks this transition is driven by the neurons with low firing rates; i.e., with larger numbers of recurrent inputs. These neurons are the first to become bursty as *J*_0_ increases (see S6 Text).

**Fig 12.**
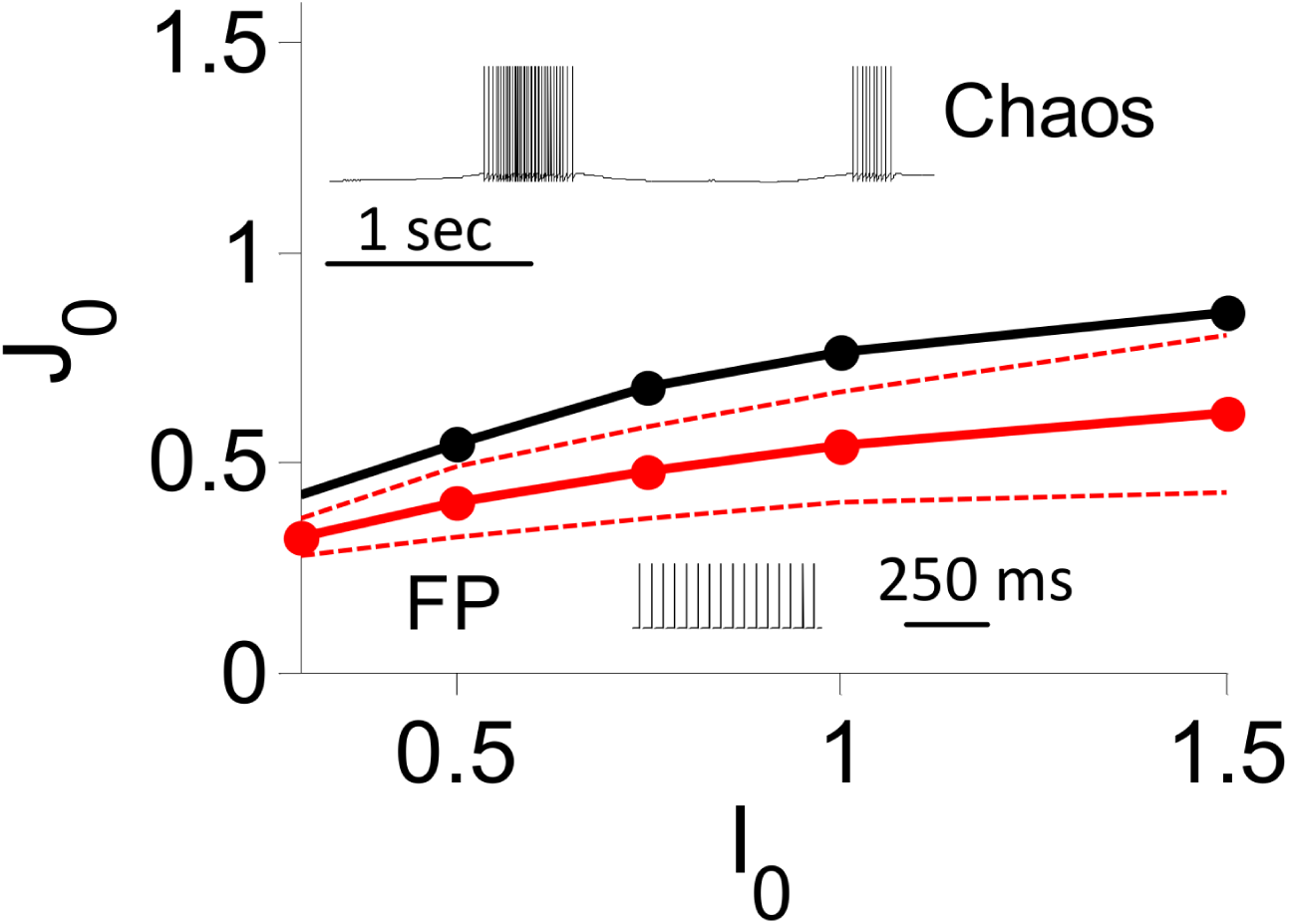
Phase diagram of the inhibitory LIF rate model. All the results are from numerical simulations with *N* = 40,000, *K* = 800. Black: zero-crossing of the maximum Lyapunov exponent, Λ. The fraction of networks for which the dynamics converge to a fixed point is 50%. 5% and 95% on the solid, top-dashed and bottom-dashed red lines respectively. Insets: *I*_0_ = 0.3. Voltage traces of a neuron in the inhibitory LIF spiking model for *J*_0_ = 2 (top inset), 0.3 (bottom inset) and *τ_syn_*=100 ms.

In Fig.13A we plot the bifurcation diagram of the model as obtained in the numerical solution of the DMFT equations (black line) and as calculated in simulations of the rate model (blue dots) and of the spiking network with *τ_syn_* = 25 ms (red × ’s) and *τ_syn_* = 7.5 ms (green × ’s). The rate model simulations are in good agreement with DMFT for 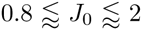. For larger *J*_0_ the discrepancy becomes significant and increases with *J*_0_. This is because of finite *K* effects that grow stronger as *J*_0_ increases as shown in the right inset in Fig.13A, for *J*_0_ = 3 (blue) and *J*_0_ = 4 (red). Fig. 13A also shows that, as discussed above, the amplitude of the PACs obtained in simulations of the LIF rate and spiking networks are barely different provided the synaptic time constant is sufficiently large.

**Fig 13.**
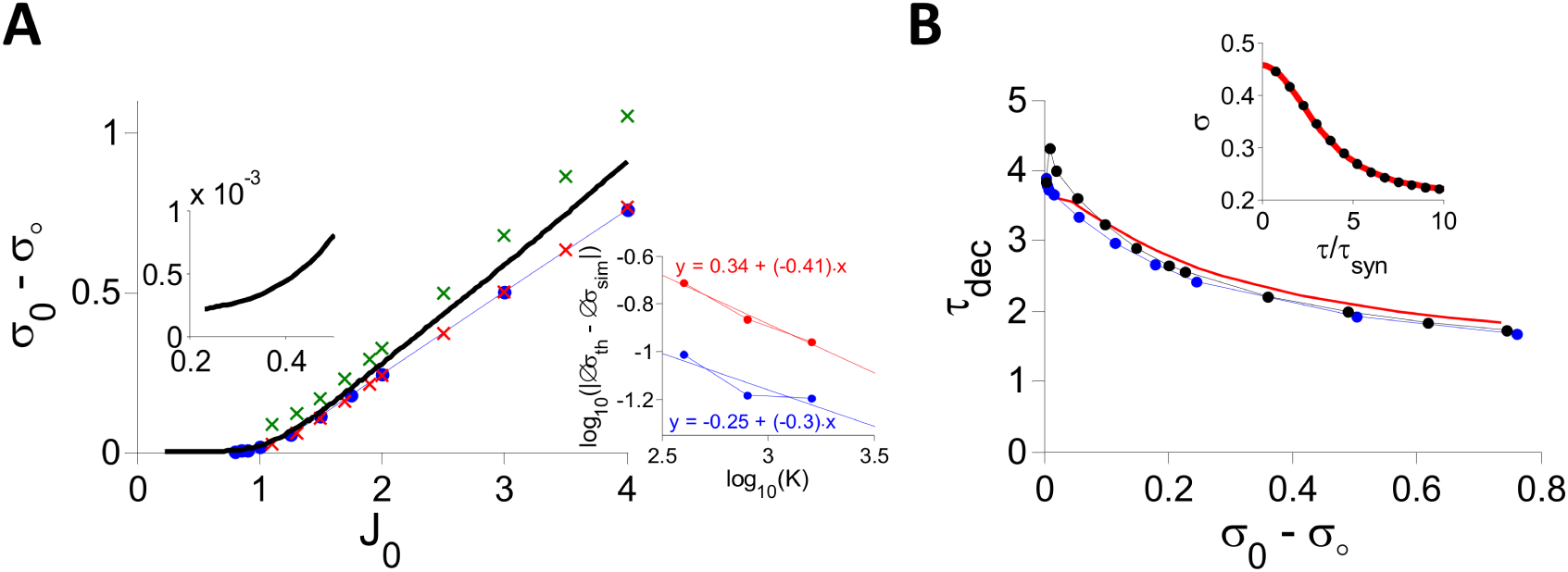
DMFT vs. numerical simulations in the one-population LIF rate model. All simulation results depicted here were obtained in networks with *N* = 40,000, *K* = 800, *I*_0_ = 0.3. A: The PAC amplitude, *σ*_0_ − *σ*_*∞*_, vs. inhibitory coupling, *J*_0_. Black: DMFT. Blue dots: Simulations of the rate model. Red *×*’s: Simulations of the spiking network with *τ_syn_* = 25 ms. Green *×*: Spiking network with *τ_syn_* = 7.5 ms. Right inset: The difference between PAC amplitudes obtained in simulations (Δ*σ_sim_*) and DMFT (Δ*σ_th_*) plotted against *K* (in log scale) for *J*_0_ = 3 (blue) and *J*_0_ = 4(red). Left inset: Closeup (*J*_0_ ∈ [0.2 0.5]) of the numerical solution of the DMFT equations. B: PACs were fitted as explained in the text to estimate *τ_dec_*. The estimated decorrelation time, *τ_dec_*, is plotted vs. the amplitude of the PAC for the rate (blue), spiking (black) networks and DMFT (red). Inset: The PAC in the rate model for *J*_0_ = 2 (black dots: simulation; red line: fit).

Fig. 13B shows the relation between the decorrelation time, *τ_dec_* and the PAC amplitude. To get these results, simulations of the rate and the spiking networks were performed for *J*_0_ ∈ [0.8, 3.5] and *τ_dec_* was estimated by fitting the PACs with the function *A ·* [cosh(*τ/τ_dec_*)]^-1^. We also solved the DMFT equations for the same values of *J*_0_ and computed the PAC that we fitted to the same function. The results from the simulations (rate model: blue; spiking network: black) and DMFT (red) agree fairly well. Note that *τ_dec_* decreases more slowly as *σ*_0_ *- σ*_∞_ increases than in the models with a sigmoid or threshold-linear transfer function (compare to Figs. 4C and 7C).

Finally, according to the DMFT the fixed point should be always unstable since for the LIF transfer function the elements of the stability matrix always have an infinite variance or, equivalently, the integral in Eq. (5) always diverges. This can be seen in the close-up in the left inset of Fig.13A, indicating that the PAC amplitude is non-zero for small *J*_0_ and that it approaches 0 very slowly as *J*_0_ decreases. By contrast, in numerical simulations in the same range of *J*_0_, the dynamics are not chaotic for most of the realizations of the network: they converge to a fixed point, as shown in Fig.12. The explanation for this difference is as for the rate model with threshold power-law transfer function with γ *<* 1*/*2 (see above).

## Two asynchronous chaos mechanisms in excitatory-inhibitory recurrent networks

We now consider EI spiking networks with recurrevt feedback interactions between the two populations.The synaptic strengths and time constants are 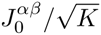 and *τ _α β_* (*α, β* ∈ {*E, I*}). Assuming slow synapses, the dynamics can be reduced to four sets of equations for the four types of synaptic inputs 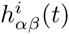 (*Materials and Methods*, Eq. (17)). The DMFT yields self-consistent equations for the statistics of these inputs. These equations can be analyzed straightforwardly for the fixed point state. In contrast to purely inhibitory networks where the fixed point loses stability only via a bifurcation to chaos, it can now also lose stability via a Hopf bifurcation. This depends on the synaptic time constants. When this happens the network develops synchronous oscillations which break the balance of excitation and inhibition (the oscillation amplitude diverges for large *K*).

We focus here on instabilities which lead to chaos. Their locations in the 6 dimensional parameter space (4 synaptic strengths, 2 external inputs) of the model can be derived for a general transfer function (Eqs. (54)-(55)). Differential equations for the PAC functions, *σ _α β_*(*τ*), can also be derived in the chaotic regime. However, general analytical characterization of their solutions is particularly difficult. Leaving such study for future work, we mostly focus below on numerical simulations. Our key result is that in EI networks asynchronous chaos emerges in two ways, one driven by I-I interactions (II mechanism) and the other by the EIE loop (EIE mechanism).

### EI network with threshold-linear transfer function

We first study a EI network in which all the neuronal transfer functions are threshold-linear. Fig. 14 plots for different *K* the phase diagram of the DMFT of this model in the 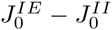 parameter space, when 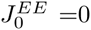 and *I*_*E*_=*I*_*I*_ =1, 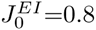.(The phase-diagram for a non-zero of value 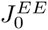, 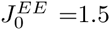, is plotted and briefly discussed in S7 Text). On the lines, the dynamics bifurcate from fixed point (below the lines) to chaos (above). As 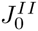 decreases the lines go to infinity. Numerical simulations indicate the maximum Lyapunov exponent changes sign very close to these lines (compare red line and red dots) in good agreement with DMFT. For any finite *K*, the instability line exhibits a re-entrance, crossing the 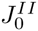-axis at 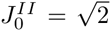, where the instability occurs in a purely inhibitory network; in this respect, the limit *K* → ∞ is singular. Solving the self-consistent equations for the average firing rates, *r*^*E*^ and *r*^*I*^, one finds that the two populations can have a comparable firing rate for large 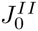 when 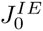 is not too large. As 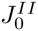 becomes small, the activity in the E population becomes much lower than in the I population. In fact, for *K* → ∞, *r*^*E*^ vanishes on the line 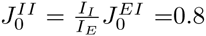 and is zero for 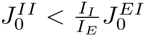 (white region in Fig.14). In other words, in the latter case, inhibition is not balanced by excitation in the E population.

**Fig 14.**
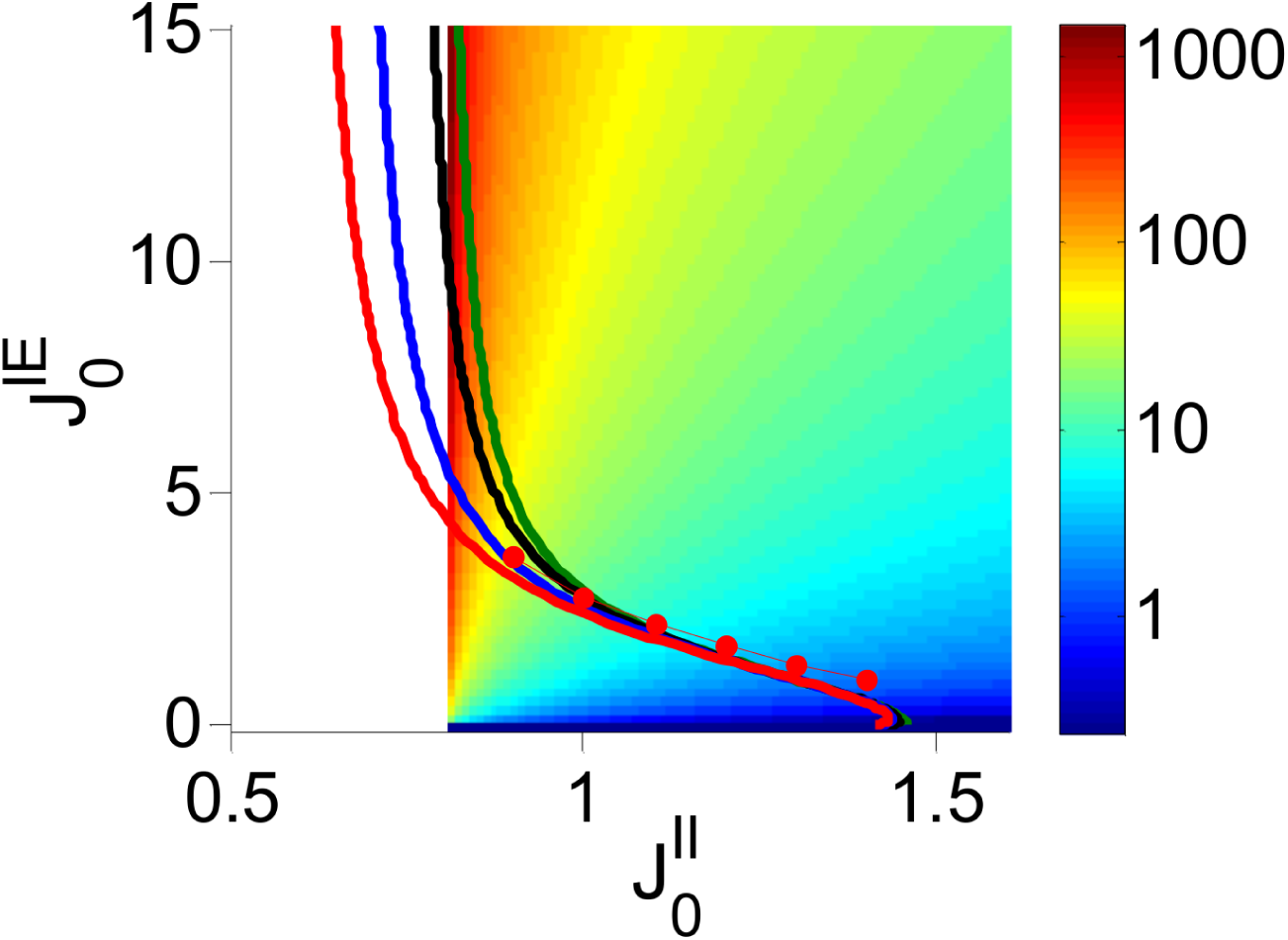
The phase diagram of the two-population rate model with threshold-linear transfer function. 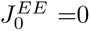, 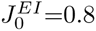. The bifurcation lines predicted by the DMFT are plotted in the 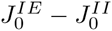 parameter space for *K* = 400 (red), 10 ^3^ (blue), 10 ^4^ (black), and *K →* ∞ (green). Red dots: Zero-crossing of the largest Lyapunov exponent (average over 5 network realizations) in numerical simulations for *K* = 400. Color code: Ratio of the population average firing rate of the two populations (I/E) in *log* scale (right).White region: The activity of the E population is very small for finite *K* and goes to zero in the limit *K* → ∞. The boundary to the right of that region is given by: 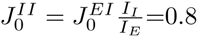.

As shown above, in the single inhibition population case with threshold-linear transfer functions the transition to chaos occurs at 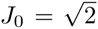 Fig. 14 shows that in the two population network the chaotic regime extends below 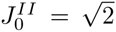. This suggests that the EIE loop can also play the key role in the emergence of chaos. To assess further the role of the II and of the EIE interactions in generating chaotic activity, we simulated the network for different values of 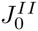 and *τ _α β_*. Traces of the synaptic inputs are displayed in Fig 15 for large (panel A) and small (panel B)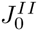. The gray traces correspond to the case where all time constants are equal (10 ms, reference case). Multiplying *τ_IE_* by 10 (black) slows down the fluctuations in all inputs when 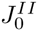 is small, but when 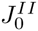 is large this happens only for *h*_*I E*_. By contrast, dividing *τ_II_* by 10 (purple) has very little effect when is small but the fluctuations of all the inputs are substantially faster when 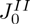 is large.

**Fig 15.**
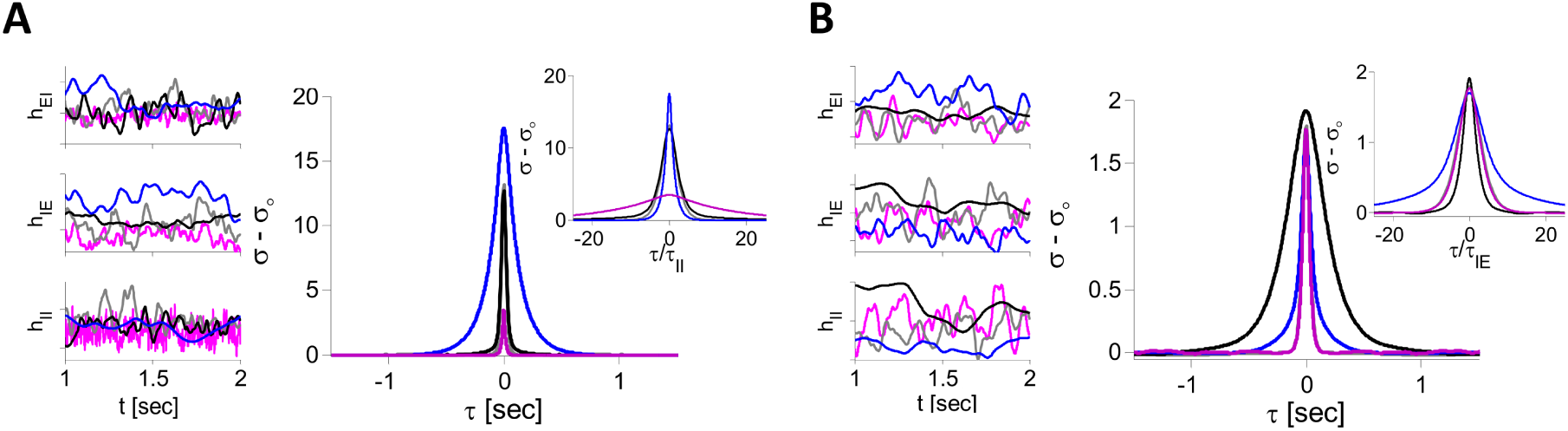
The two mechanisms for asynchronous chaos in the two-population rate model with threshold-linear transfer function. Simulations were performed for *N*_*E*_ = *N*_*I*_ = 8000, *K* = 400, *I*_*E*_ = *I*_*I*_ = 1, 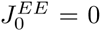, 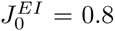. A: II mechanism for 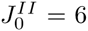, 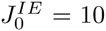. Left panels: Examples of traces of excitatory (*h*_*IE*_) and inhibitory inputs (*h*_*EI*_, *h*_*II*_) into one neuron. Right: PAC of the net inputs to the E neurons. Gray: *τ_IE_* = *τ_EI_* = *τ_II_* = 10 ms; Black: *τ_IE_* = 100 ms, *τ_EI_* = *τ_II_* = 10 ms; Blue: *τ_II_* = 100 ms, *τ_IE_* = *τ_EI_* = 10 ms; Purple: *τ_II_* = 1 ms, *τ_EI_* = *τ_IE_* = 10 ms. Inset: All PACs plotted vs. *τ /τ_II_*. B: EIE mechanism for 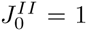, 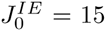. Other parameters are as in A. Inset: All PACs plotted vs. *τ /τ_IE_*.

Fig. 15 also demonstrates the effect of changing *τ _α β_* on the PAC of the net inputs to the E neurons, 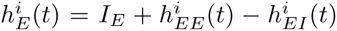 (corresponding results for the I population are shown in S8 Text). The PAC in the reference case is plotted in gray. For large 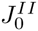, a ten-fold *increase* in *τ_II_* causes the PAC width to become ten times larger and the PAC amplitude increases (Fig 15A, blue; see also inset). For a ten-fold *decrease* in *τ_II_* (purple) compared to reference, the width of the PAC is smaller but by a smaller factor whereas its amplitude is greatly reduced. By contrast, a ten-fold increase in *τ_IE_* has no noticeable effect, either on the width or on the amplitude of the PAC (black). Fig. 15B plots the PAC of the total input to the E population for small 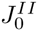. Here, decreasing *τ_II_* by a factor of 10 (purple line) weakly affects the width as well as the amplitude of the PAC. In contrast, a ten-fold increase of *τ_IE_* (black) widens the PAC by a comparable factor (see also inset). A similar widening occurs if *τ_EI_* is increased ten-fold (see S8 Text).

This phenomenology can be understood as follows. In the large 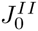 regime, the II interactions play the key role in the generation of chaos. Therefore, the time scale of the fluctuations in the activity of the I neurons is essentially determined by *τ_II_*. Thus if the latter is 10 times larger than reference, the I inputs to the E neurons are slowed down by the same factor. At the same time, the filtering effect of the EI synapses becomes weaker and thus the amplitude of the PAC of the net input in the E neurons increases. The effect of decreasing *τ_II_* stems from the filtering effect of the EI synapses which is now stronger than in the reference case. Finally, changing *τ_IE_* has no noticeable effect since the fluctuations are generated by the II interactions. By contrast, when 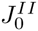 is small, II interactions are not sufficient to generate chaotic fluctuations. In this regime, the EIE loop drives these fluctuations if 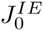 is sufficiently large. That is why the time scale of the activity fluctuations depends primarily on *τ_IE_* and to a much smaller extent on *τ_II_*.

These results point to the existence of two mechanisms for chaos emergence in two population networks; they differ by the type of the dominant interactions (EIE or II) and therefore on the synaptic time constants which settle the time scale of the activity fluctuations. Another difference is that in the EIE mechanism, the E population is always significantly less active than the I population. This is not the case in the II mechanism.

### Two-population spiking LIF network

We ran a similar analysis for LIF networks. Fig.s 16A,C plot the PACs of 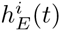 for the LIF spiking and rate models (PACs of 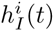 are shown in S9 Text). In all panels 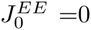, 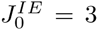, 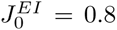 and *τ_EI_* =3 ms. For 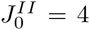 (Fig 16A), increasing *τ_II_* slows down the fluctuations. By contrast, changing *τ_IE_* only has a very mild effect (S10 Text). This is because the fluctuations are essentially driven by the II interactions. For 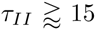, the fluctuation statistics are quantitatively similar in the spiking and the rate models: in both, the decorrelation time, *τ_dec_* ≧ 2*τ_II_* (Fig 16A, inset). Moreover, simulations indicate that the dynamics of the rate model are chaotic (λ ≈ 1.7*/τ_II_*). The trace in Fig 16B shows that with large *τ_II_* (=100 ms) the spiking pattern is bursty. The membrane potential between between bursts exhibit slow fluctuations because they are generated by the slow II connections.

**Fig 16.**
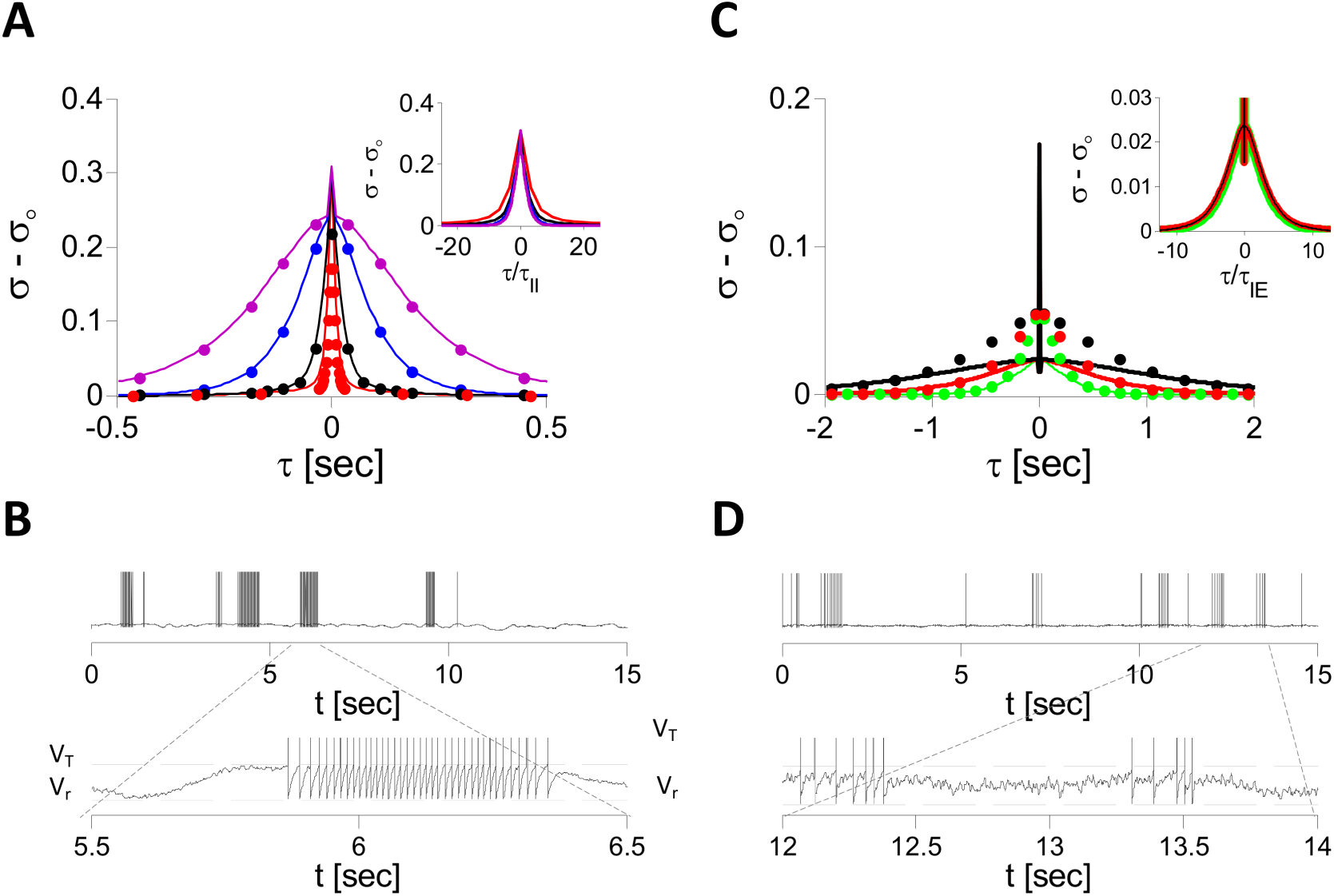
The two mechanisms for asynchronous chaos in two-population LIF spiking and rate networks. Simulations were performed with *N*_*E*_ = *N*_*I*_ = 16000, *K* = 400,*I*_*E*_ = 0.2, *I*_*I*_ = 0.1, 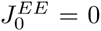, 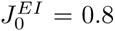, 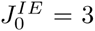 A: II mechanism. PACs of the net inputs in E neurons are plotted for, *τ_IE_* = 100 ms, *τ_EI_* = 3 ms and *τ_II_* = 3, (red), 10 (black), 40 (blue) and 100 ms (purple). Solid line: Spiking model. Dots: Rate model. Inset: All PACs (spiking network) are plotted vs. *τ /τ_II_*. B: Voltage of one E neuron for parameters as in A, purple. C: EIE mechanism. PACs of the net inputs in E neurons are plotted for 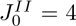, *τ_EI_* = *τ_II_* = 3 ms and *τ_IE_* = 100, (green), 200 (red) and 400 ms (black). Solid line: Spiking model. Dots: Rate model. Inset: All PACs (spiking network) are plotted vs. *τ /τ_IE_*. D: Voltage of one E neuron in the spiking network with parameters as in C, green.

Fig. 16C plots the PACs of 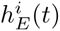 for 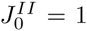. Here also, the LIF rate model operates in a chaotic regime (λ ≈ 120*s*^-1^). In the spiking model the PACs exhibit a slow time scale but also a fast one (the sharp peak around *τ* = 0). These correspond to the slow and fast fluctuations observable in the voltage traces in Fig 16D. Increasing *τ_IE_* while keeping *τ_EI_* = *τ_II_* = 3 msec has a substantial effect on the slow component but hardly a?ects the fast component. When plotted vs. *τ /τ_IE_*, the slow components of the PACs all collapse onto the same curve (Fig 16C, inset). This indicates that the EIE loop is essential in generating the slow, but not the fast, fluctuations. Fitting this slow component with the function *A ·* [cosh(*τ /τ_dec_*)]^-1^ yields *τ_dec_* ≈ 2.4*τ_IE_*. Furthermore, increasing *τ_II_* suppresses the fast fluctuations and amplifies the slow ones. These two effects saturate simultaneously when *τ_II_* ≈ 10 ms (S11 Text). Thus, it can be inferred that fast fluctuations are mostly generated by II interactions. Their amplitude is suppressed as *τ_II_* is increased because they become more filtered. Concomitantly, the slow fluctuations become amplified. This is because fast fluctuations smooth the effective transfer function of the E neurons in the low firing rate regime. Thus, their suppression increases the gain of this transfer function. This explains the quantitative differences between the PACs in the spiking and the rate LIF network when II synapses are fast and why these differences are lessened as *τ_II_* increases (S11 Text).

In the simulations reported in Fig 16 there is no recurrent excitation in the E population 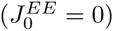. Moreover, all the excitatory synapses to the I population are slow. Both assumptions were made to reduce the number of parameters in order to simplify the analysis. However, in cortical networks in general, fast (AMPA) and slow (NMDA) excitation coexist (in fact AMPA synapses are required to open the NMDA receptors). Moreover, recurrent excitation is thought to be in general substantial (see however [49]). Results depicted in S12 Text show that the EIE loop can induce slow rate fluctuations in our network when it combines slow and fast excitatory synapses and when substantial recurrent excitation is present in the E population.

## Discussion

Networks of neurons operating in the so-called balanced regime exhibit spiking activity with strong temporal variability and spatial heterogeneity. Previous theoretical studies have investigated this regime assuming that excitatory and inhibitory synapses are sufficiently fast compared to the neuronal dynamics. The nature of the balanced state is now fairly well understood in this case. By contrast, here we focused on networks in which some of the synapses are slow. To study the dynamics in these networks, we reduced them to a rate dynamics that we investigated by combining Dynamical Mean-Field Theory and simulations. Our key result is that when synaptic interactions are sufficiently strong and slow, chaotic fluctuations on the time scales of the synaptic dynamics emerge *naturally* from the network collective behavior. Moreover, the nature of the transition to chaos and the behavior in the chaotic regime are determined only by the neuronal *f-I* curve and not by the details of the spike-generation mechanism.

We identified two mechanisms for the emergence of asynchronous chaos in EI neuronal networks. One mechanism relies on II interactions whereas in the other the EIE feedback loop plays the key role. These mechanisms hold in rate models (Eq. (3)) as well as in LIF spiking networks. By computing the maximum Lyapunov exponent, we provided direct evidence that in rate models these states are indeed chaotic. For LIF spiking networks, we argued that when the synapses are sufficiently slow, the observed activity fluctuations are chaotic since their statistics are quantitatively similar to those observed in the corresponding rate model. This similarity persists for synaptic time constants as small as the membrane time constant. This is in agreement with [33–35] which relied on numerical integration of the LIF model to compute the Lyapunov spectra of networks of various sizes and increasing synaptic time constants. They found that the LIF dynamics are chaotic *only* if the synapses are sufficiently slow.

In these two mechanisms, the dynamics of the synaptic currents play the key role whereas dependence on the intrinsic properties of the neurons only occurs via their nonlinear instantaneous input-output transfer function. Since the synaptic currents are filtered versions of the neuronal spike trains, and that the temporal fluctuations of the activity occur on the time scales of the synaptic currents, it is natural to qualify the dynamical regime as *rate* chaos. Although the features of the bifurcation to chaos may depend on the shape of the transfer function, as we have shown, the qualitative features of the chaotic state are very general, provided that the synaptic currents are sufficiently slow. Rate chaos is therefore a generic property of networks of spiking neurons operating in the balanced regime. We show in S3 Text that rate chaos occurs also in networks of *non-leaky* integrate-and-fire spiking neurons. In that case, the statistics of the fluctuations are similar to those of the model in Eq. (3) with a threshold-linear transfer function. We also found rate chaos in biophysically more realistic network models in which the dynamics of the neurons and of the synapses are conductance-based (results not shown). In these cases, the dynamics of the synaptic conductances give rise to the chaotic fluctuations.

Quantitative mappings from spiking to rate models have been derived for networks in stationary asynchronous non chaotic states [39] or responding to external fluctuating inputs [48]. Spiking dynamics also share *qualitative* similarities with rate models for networks operating in synchronous states [9–11, 38, 39]. To our knowledge, the current study is the first to report a *quantitative* correspondance between spiking and rate model operating in chaotic states.

The SCS model [19] has been widely used to explore the physiological [22, 50] and computational significance of chaos in neuronal networks. Recent works have shown that because of the richness of its chaotic dynamics, the SCS model has remarkable learning capabilities [15–18]. Our work paves the way for an extension of these results to networks of spiking neurons with a connectivity satisfying Dale’s law, which are biologically more realistic than the SCS model.

Another interesting implication of our work is in the field of random matrices. Given a *dense* NxN random matrix, **A**, with i.i.d element with zero mean and finite standard deviation (SD), in the large *N* limit, the eigenvalue of 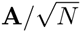 with the largest real part is real, and it is equal to SD [51, 52] (more generally, the eigenvalues of 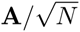 are uniformly distributed within a disk of radius SD centered at the origin [51, 52]). Several results regarding the spectra (bulk and outliers) of dense random matrices with structures reflecting Dale’s law have been derived recently [53–55]. Less is known when the matrices are sparse. A byproduct of our approach are two conjectures for the maximal eigenvalue of such *sparse* random matrices, namely Eqs. (7) and (62) that we verified numerically.

Neuronal spiking statistics (e.g., firing rate, spike counts, inter-spike intervals) exhibit a very broad range of time scales during spontaneous or sensory evoked activity in-vivo (see e.g [56, 57]). Fluctuations on time scales larger than several 100s of millisecond can be accounted for by neuromodulation which changes the global excitability of the cortical network or changes in behavioral state. Very fast fluctuations are naturally explained in the framework of the standard model of balance of excitation and inhibition [28–30]. By contrast, it is unclear how to explain modulations in the intermediate temporal range of a few 10s to several 100s of milliseconds. In fact, the *standard* framework of balanced networks predicts that fluctuations on this time scale are actively suppressed because the network state is very stable. Our work extends this framework and shows two mechanisms by which modulations in this range can occur. In the II mechanism, inhibitory synapses must be strong and slower than 10 - 20 ms. GABA_A_ inhibition may be too fast for this [58] (see however [59]), but GABA_B_ [60] are sufficiently slow.In contrast, the EIE mechanism is achieved when inhibition in fast. It requires slow recurrent excitation to inhibitory neurons, with a time constant of a few to several tens of ms, as is typically the case for NMDA receptors (see e.g [61–63]). Hence, the combination of GABA_A_ and NMDA synapses can generate chaotic dynamics in the cortex and fluctuations in activity on a time scale of several tens to a few hundreds of ms.

**Note added in production:** Following a request from the editors after formal acceptance of our article, we note that a recent paper [64] claims that spiking networks with instantaneous delayed synapses exhibit an asynchronous state similar to the chaotic state of the SCS model. However, this claim is incorrect and has been shown to rely on flawed analysis [65].

## Materials and Methods

### Models

#### Two population leaky integrate-and-fire spiking network

The two population network of leaky integrate-and-fire (LIF) neurons considered in this work consists of *N*_*E*_ excitatory (E) and *N*_*I*_ inhibitory neurons. The subthreshold dynamics of the membrane potential 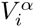, of neuron *i* in population *α* (i=1,…,*N _α_*; α, *β* ∈ {*E, I*}) obeys:

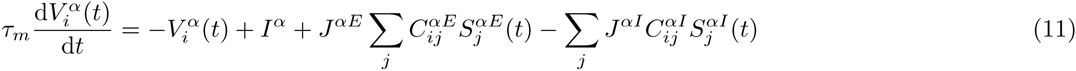

where *τ_m_* is the membrane time constant (we take *τ_m_* = 10 msec for both populations), 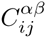 and *J*^*α β*^ are respectively the connectivity matrix and the strength of the connections between the (presynaptic) population β and (postsynaptic) population *α* and *I*^*α*^ the external feedforward input to population *α*. For simplicity we take *N*_*E*_ = *N*_*I*_ = *N*. However, all the results described in the paper are also valid when the number of neurons is different in the populations (provided both numbers are large)., The variables 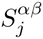, which describe the synapses connecting neuron *j* in population *β* to population *α*, follow the dynamics:

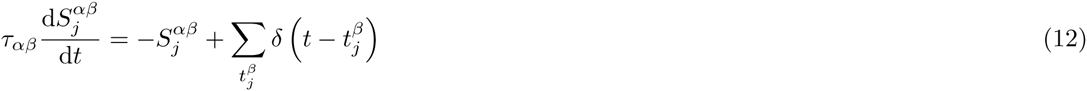

where *τ _α β_* is the synaptic time constant and the sum is over all the spikes emitted at times 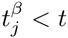.

Equations (11,12) are supplemented by a reset condition. If at time *t*_*sp*_, 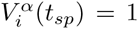, the neuron emits a spike and 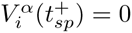. For simplicity we do not include the neuronal refractory period.

We assume that the connectivity is random with all the 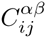 uncorrelated and such that 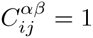 wit probability *K/N* and 0 otherwise. Hence each neuron is connected, on average, to *K* neurons from its population as well as to *K* neurons from the other population. When varying the connectivity *K* we scale the interaction strength and the feedforward inputs according to: 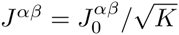 and 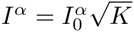 [29].

#### Network of inhibitory leaky integrate-and-fire neurons

The dynamics of the network of the one-population spiking LIF neurons considered in the first part of the paper are:

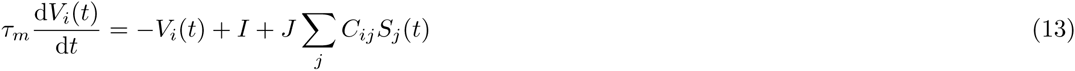

supplemented with the reset condition at threshold. The elements of the connectivity matrix, *C*_*ij*_, are uncorrelated and such that *C*_*ij*_ = 1 with probability *K/N* and 0 otherwise. All neurons are inhibitory, thus *J* < 0.

The synaptic dynamics are:

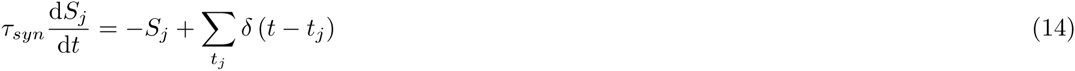

where *τ_syn_* is the synaptic time constant of the inhibition and the sum is over all the spikes emitted at times *t*_*j*_ < *t*. The interaction strength and the feedforward inputs scale with K as: 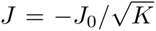 and 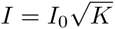 with *J*_0_ *>* 0.

#### Network of non-leaky integrate-and-fire neurons

We consider briefly this model in S3 Text. The network architecture as well as the synaptic dynamics are as above. The single neuron dynamics of non-leaky integrate-and-fire (NLIF) neurons are similar to those of LIF neurons except for the first terms on the right-hand side of Eqs. (11,13) which are now omitted.

#### Rate dynamics for spiking networks with slow synapses

If the synapses are much slower than the membrane time constant, the full dynamics of a spiking network can be approximated by the dynamics of the synapses driven by the instantaneous firing rates of the neurons, namely:

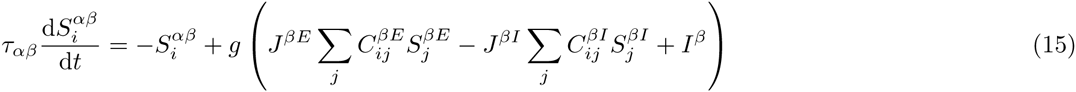

where *g*(*x*) is the transfer function of the neuron (the *f-I* curve) [20]. In particular, for the LIF networks,

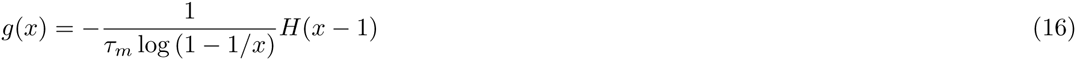

with *H*(*x*) = 1 for *x* > 0 and *H*(*x*) = 0 otherwise. For the NLIF networks, the transfer function is threshold-linear: *g*(*x*) = *xH*(*x*).

Defining 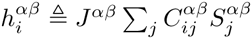, the dynamics of 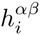 are given by

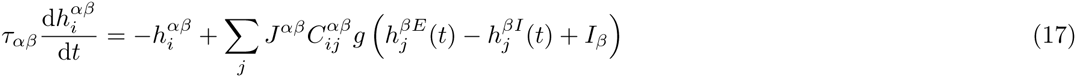

We will denote by 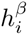 the total input into neuron *i* in population *β*: 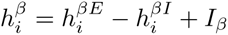. For networks comprising only one population of inhibitory spiking neurons we will drop the superscript *β* = *I* and denote this input by *h*_*i*_. The dynamics then yield:

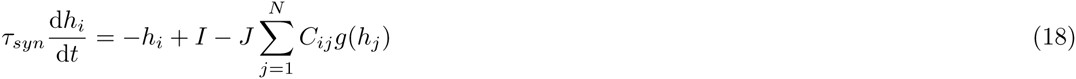

where *τ_syn_* is the inhibitory synaptic time constant.

### Dynamical Mean-Field Theory of the Single Inhibitory Population

A Dynamical Mean-Field Theory (DMFT) can be developed to investigate the rate model, Eq. (17), for a general transfer function under the assumption, 1 ≪ *K* ≪ *N*.

Here we provide a full analysis of a one-population network of inhibitory neurons whose dynamics are given in Eq. (18). We take 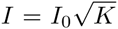 as the external input and 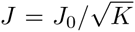 as the coupling strength In this case, a functional integral derivation shows that these dynamics can be written as:

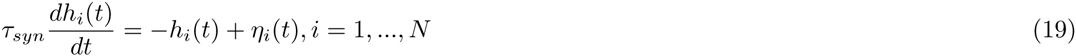

where *η_i_*(*t*) is a Gaussian noise:

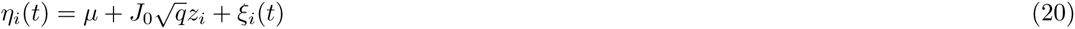

with *z*_*i*_, i.i.d Gaussian quenched variables with zero mean and unit standard deviation (SD), *ξ_i_*(*t*) are Gaussian noises with 〈*ξ_i_*(*t*)〉 _*t*_ = 0, and 〈*ξ_i_*(*t*)*ξ_j_*(*t* + *τ*)〉_*t*_ = *C*_*ξ*_(*τ*) *δ _i,j_* where 〈·〉_*t*_ stands for averaging over time. Therefore, in general, the inputs to the neurons display temporal as well as quenched fluctuations.

The self-consistent equations that determine the mean, temporal correlations and quenched fluctuations yield:

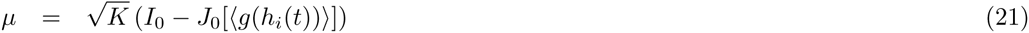

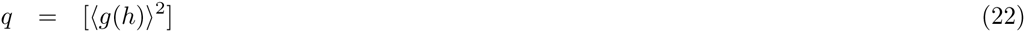

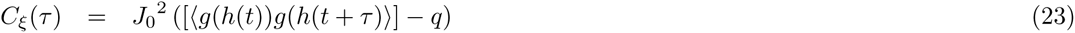

where 〈·〉and [·] stand for averaging over noise and quenched disorder, respectively. Thus the quantities *q* and *μ* obey:

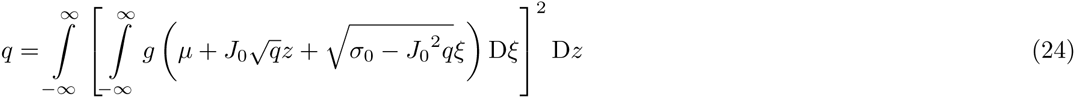

and

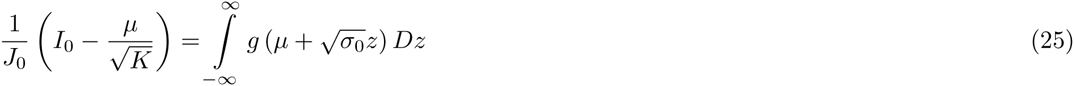

where *σ*(*τ*) = [〈*h*(*t*)*h*(*t* + *τ*〉] − *μ*^2^ is the population-averaged autocovariance (PAC) of the input to the neurons and we define: *σ*_0_ = *σ*(0) and 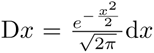 In the limit *K → ∞*, *μ* must remain finite. This implies that the population averaged firing rate, [〈*g*(*h*)〉 = *I*_0_*/J*_0_] does not depend on the specifics of the transfer function of the neurons and varies linearly with *I*_0_. This is a key outcome of the balance between the feedforward excitatory and the recurrent inhibitory inputs to the neurons.

To express *C*_*ξ*_(*τ*) in terms of *σ*, we note that the vector (*h*_*T*_(*t*)*, h*(*t* + *τ*))^*T*^ is a bivariate Gaussian, so in fact we need to calculate *E* [*g* (*μ* + *x*) *g* (*μ* + *y*)] where (*x, y*)^*T*^ has zero mean and a covariance matrix

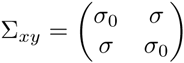

and *E*[*·*] stands for averaging over temporal noise and quenched disorder. Defining

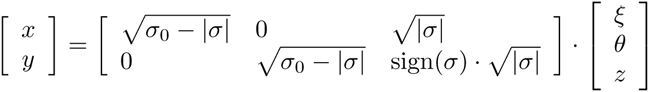

where *ξ*, *θ* and *z* are independent Gaussian variables with zero mean and unit variance yields

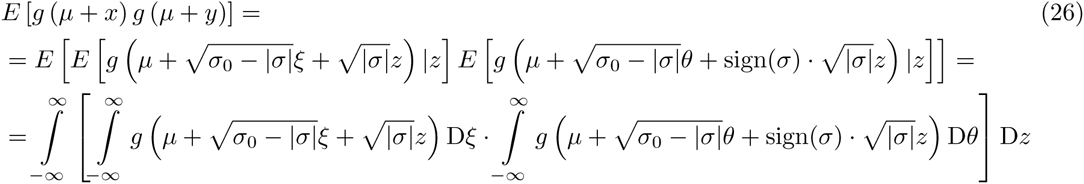

A straightforward derivation shows that *σ*(*τ*) obeys:

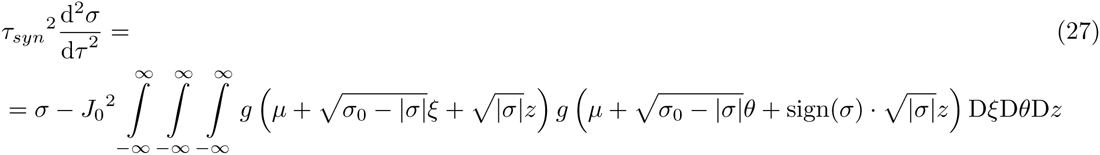

with initial conditions:

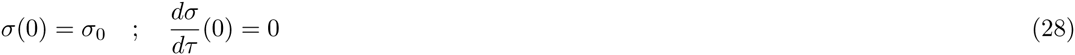

where the last condition results from *σ*(*τ*) = *σ*(*-τ*).

Equation (27) can be rewritten as:

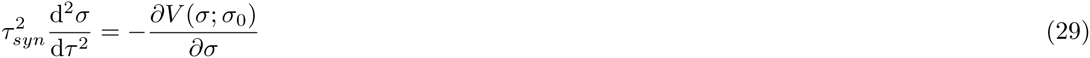

where the “potential” *V* (*σ; σ*_0_) which depends parametrically on *σ*_0_ is:

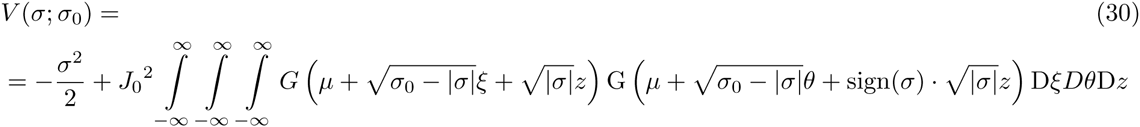

with *G*(*x*) = ∫*g*(*x*)*dx*. Note that for positive *σ* this equation yields

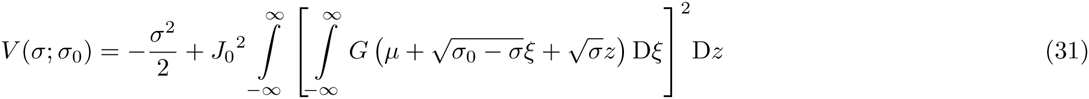

Therefore the quantity

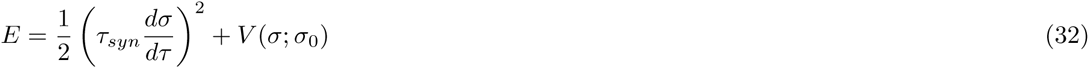

is conserved under the dynamics, Eq. (29). Hence:

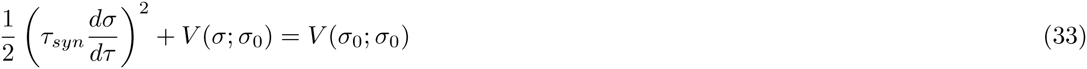

To simplify notations, we drop the parameter *σ*_0_ and denote the potential by *V* (*σ*). The first, second and third order derivatives of the potential with respect to *σ* are denoted *V*′(*σ*), *V*˝(*σ*) and *V* ‴(*σ*).

For illustrative purpose, we consider a sigmoid transfer function,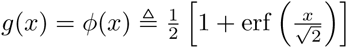.In this case we have

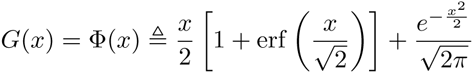

Using the identities:

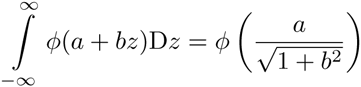

and

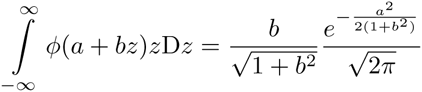

the potential *V* (*σ*) can be written as:

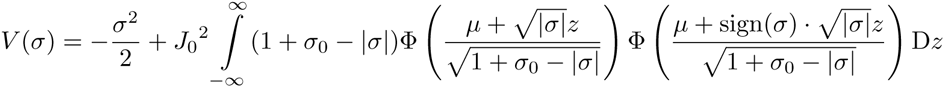

Fig. 17A_1*-*3_ plots *V* for *σ ∈* (*-σ*_0_*, σ*_0_) for *J*_0_ = 4, fixed *I*_0_ = 1 and different values of *σ*_0_. When *V ′*(*σ*_0_) *>* 0 (Fig 17*A*_1_), the solution to Eq. (29), *σ*(*τ*), decreases monotonically from *σ*_0_ to −*σ*_0_ that it reaches in finite time with a strictly negative velocity; this solution does not correspond to an autocovariance function.

**Fig 17.**
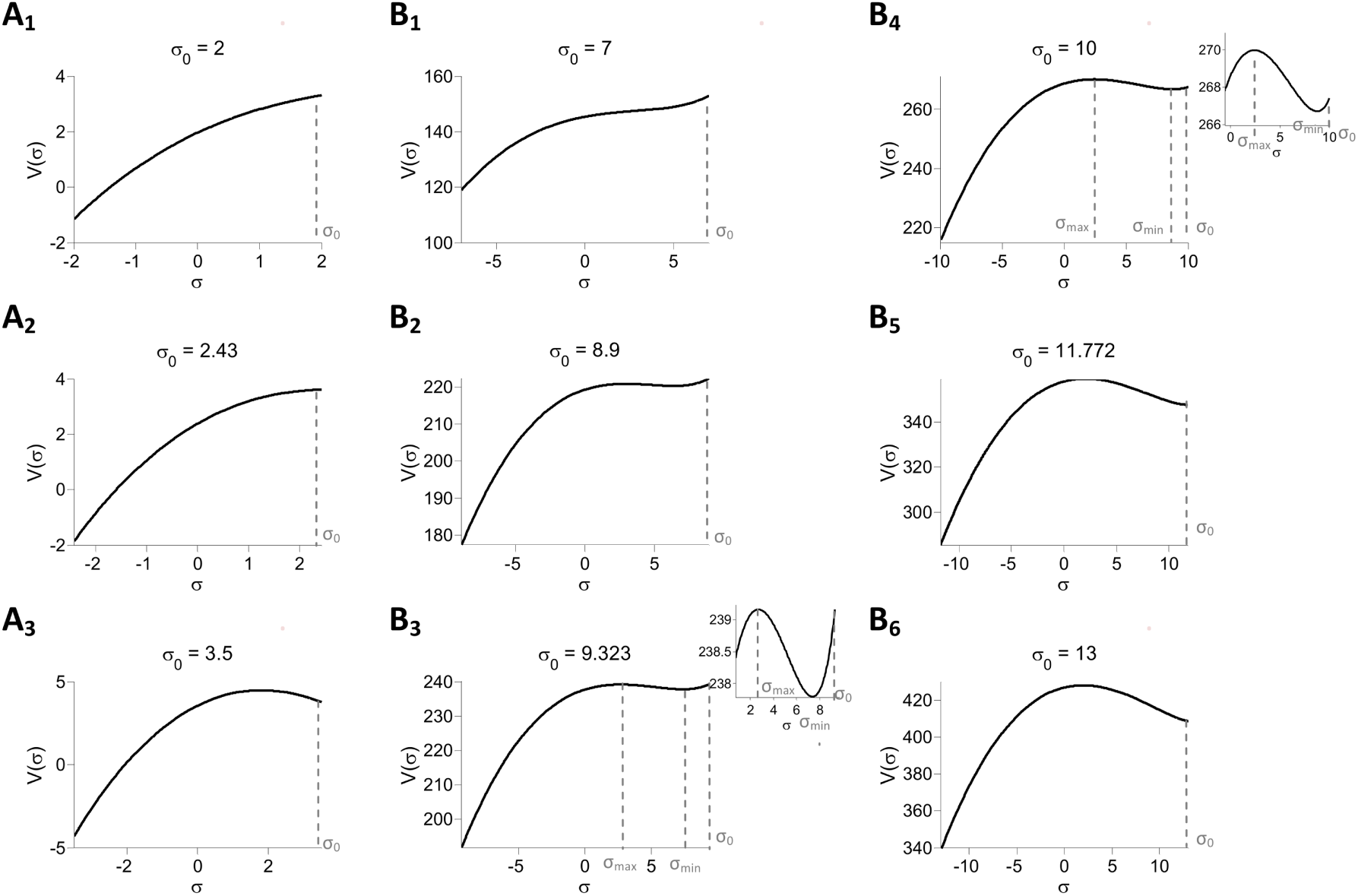
Dynamical Mean-Field Theory for the one-population inhibitory rate model with *g*(*x*) = *ϕ* (*x*). The potential, *V* (*σ, σ*_0_) is plotted for different values of *σ*_0_ as a function of *σ*. A_1*-*3_: *J*_0_ = 4 *< J_c_* (=4.995). B_1*-*5_: *J*_0_ = 15 *> J_c_*.

For *σ*_0_ such that *V*′(*σ*_0_) = 0 (Fig 17A_2_) the solution is *σ*(*τ*) = *σ*_0_. It corresponds to a fixed point of the dynamics, Eq. (18) in which all the inputs to the neurons are constant in time, 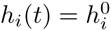, and 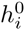 has a Gaussian distribution. Finally, for *σ*_0_ such that *V*′(*σ*_0_) *<* 0 (Fig 17A_3_), there is no solution to Eq. (33) with *σ*(0) = *σ*_0_.

Fig. 17B_1*-*3_ plots *V* for *J*_0_ = 15. For small *σ*_0_, the solution Eq. (33) does not correspond to an autocovariance function. As *σ*_0_ increases, *V* (*σ*) becomes non-monotonic in the vicinity of *σ* = *σ*_0_ with local maxima and minima at *σ* = *σ_max_* and *σ* = *σ_min_*, respectively (Fig 17B_2_). However, here also the solution for *σ*(*τ*) does not correspond to an autocovariance because *σ*_0_ is the global maximum in the range *σ ∈* [*-σ*_0_*, σ*_0_]. For 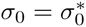, such that 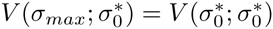 (Fig 17B_3_) an acceptable solution appears, in which *σ* decays monotonically from 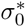 and converges to *σ_max_* as *τ* → ∞ i.e. *σ_max_* = *σ _∞_*. This solution corresponds to a chaotic state of the network. If *σ*_0_ is further increased beyond 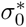, *V* (*σ_max_, σ*_0_) > *V* (*σ*_0_) (Fig 17B_4_), and the solution exhibits oscillations around *σ_min_*. For *σ*_0_ ≈ 11.77, *V ^'^*(*σ*_0_)=0, and the solution corresponds to a fixed point (Fig 17B_5_). Finally, for *σ*_0_ larger, *V*′(*σ*_0_) is negative (Fig 17B_6_) and there is no solution to Eq. (18) with *σ*(0) = *σ*_0_.

A bifurcation between these behaviors occurs at some critical value, *J*_*c*_, such that for *J*_0_ *< J_c_* the self-consistent solutions of Eq. (29) are either oscillatory or constant as a function of *τ*, whereas for *J*_0_ *> J_c_* they are either oscillatory or decay monotonically. A stability analysis of these different solutions is beyond the scope of this paper; instead, we rely on numerical simulations of the full dynamics. They indicate that the network dynamics always reach a fixed point for sufficiently small *J*_0_. For sufficiently large *J*_0_ the fixed point is unstable and the network settles in a state in which *σ*(*τ*) decays monotonically with *τ*. Simulations also show that the maximum Lyapunov exponent in these cases is positive (see below); i.e. the network is in a chaotic state. For values of *J*_0_ in between these two regimes, the network displays oscillatory patterns of activity. However, for increasing network sizes, *N*, the range of *J*_0_ in which oscillations are observed vanishes (not shown). Therefore for large *N* the bifurcation between a fixed point and chaos occurs abruptly at some critical value *J*_*c*_. A similar phenomenology occurs for other non-linear positive monotonically increasing transfer functions.

In summary, for a fixed feedforward input, *I*_0_, there are two regimes in the large *N* limit: for *J*_0_ *< J_c_*: the stable state is a fixed point. The distribution of the inputs to the neurons is a Gaussian whose mean, *μ*, and variance, *σ* are determined by the self-consistent mean-field equations:

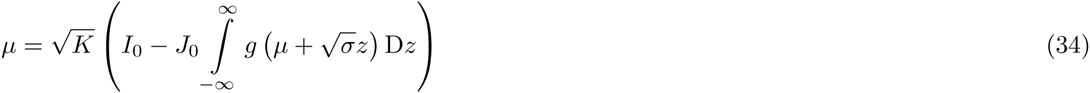

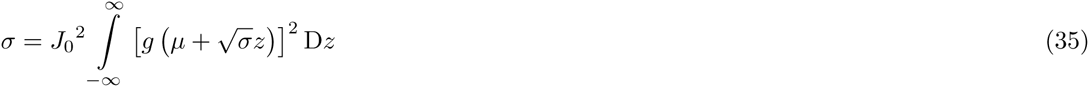

For a transfer function, *g*(*x*), which is zero when *x* is smaller than some threshold *T* (functions without threshold correspond to *T* = −∞), the distribution of the neuronal firing rates, *r*_*i*_, in this state is given by:

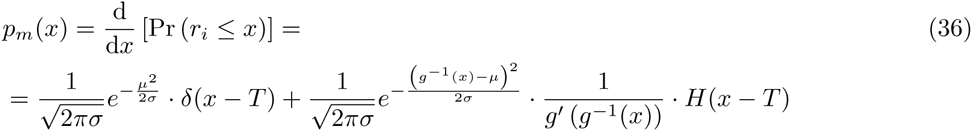

2) for *J*_0_ *> J_c_*: the stable state is chaotic. The distribution of time average inputs is Gaussian with mean *μ* and variance 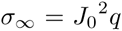 and the autocovariance of the inputs is determined by Eq. (29) which depends on *σ*_0_. The quantities *μ*, *σ*_0_ and *σ _∞_* are determined by the self-consistent equations:

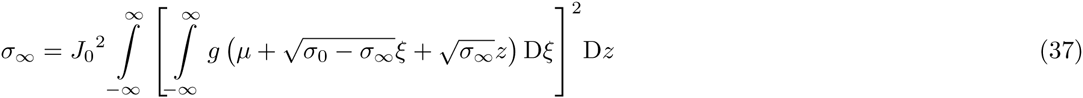

and

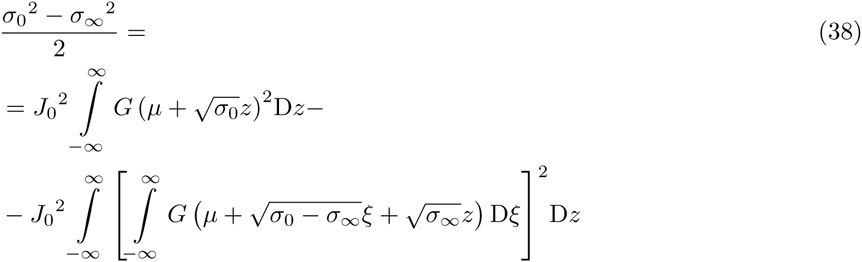

together with Eq. (25).

### Two-population networks

#### Self-consistent DMFT equations

A DMFT approach can also be developed to investigate the dynamics of the two population network model, Eq. (17). To that end, the last term in Eq. (17) is written as a Gaussian random process with mean *μ ^α β^* and autocorrelation function *C*^*α β*^ (*τ*) and derives the self-consistent equations that these quantities satisfy. The quantity *μ ^α β^* is therefore

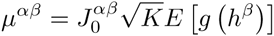

where:

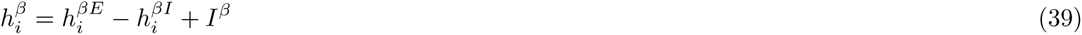

is the net input to neuron *i* in population *β*.

The synaptic inputs 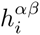 is also a Gaussian random process. We denote its mean over time and over all the neurons in population *α* by *μ ^α^ ^β^* = [*E h^α^ ^β^* (*t*)] and its PAC by *σ ^α^ ^β^* (*τ*) = *E* [*h*^*α β*^ (*t*)*h*^*α β*^ (*t* + *τ*)]*-*(*μ^α^ ^β^*)^2^. Taking 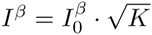 we can write the mean of 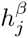 as

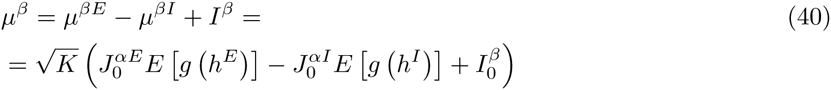

The PAC of 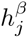 then reads:

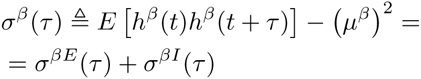

We can now write the balance condition in the large *K* limit:

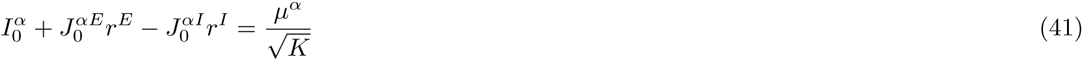

where

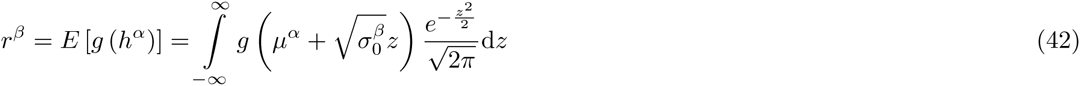

is the neuronal firing rate averaged over cells in population *α*. Here, 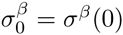.

We can also express *C ^α^ ^β^*(*τ*) in terms of *σ ^α^* (*τ*) as:

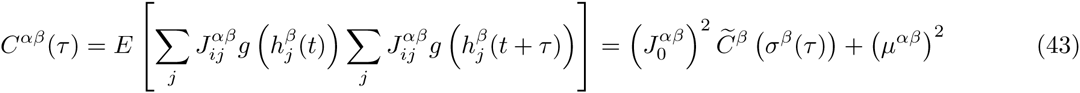

where:

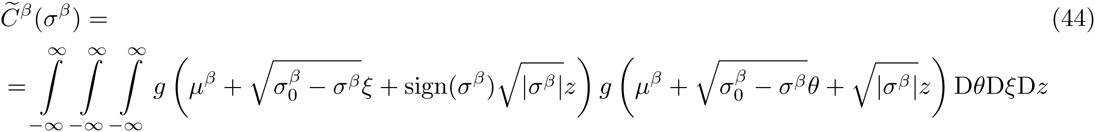

Let us denote by Δ^*α β*^(*τ*) the autocorrelation of *h*^*α β*^(*t*). We can express the relation between *C*^*α β*^(*τ*) and Δ^*α β*^(*τ*) by their Fourier transforms as Δ^*α β*^(*ω*) = *H*(*ω*)*H*^*\**^(*ω*)*C*^*α β*^(*ω*), where *H*(*ω*) = 1*/*(1 + *iτ ^α^ ^β^ ω*). Transforming back to the time domain yields:

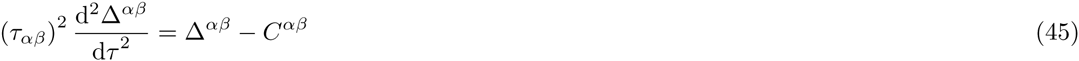

since 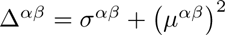 we get:

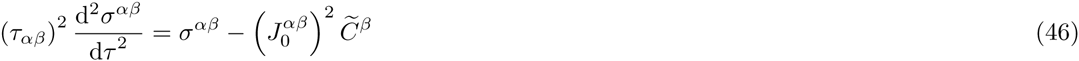

Thus we get a set of self-consistent equations for the four PACs *σ ^α β^*. The relevant soutions have to satisfy the four boundary conditions:

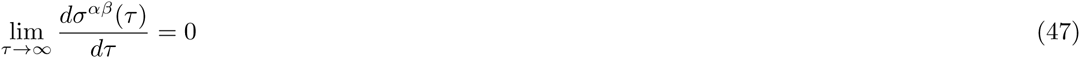

In general, these dynamical equations cannot be written like those of a particle in some potential. This makes the study of their solutions substantially more difficult than in the one population case.

#### Separation of time scales

A potential function can be written for the DMFT if the time scale of one type of synapses is substantially larger than the others, which makes it possible to consider the latter as instantaneous. We carry out this analysis below assuming *τ_IE_* ≫ *τ_EI_, τ_EE_, τ_II_*.

Setting all the synapses except those from E neurons to I neurons to be instantaneous implies that except for *σ_IE_* one has:

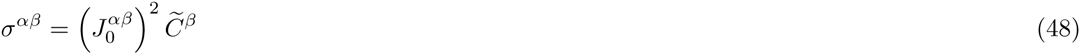

where 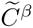 is defined in Eq. (44). Since *τ_IE_* is now the only time scale we can take *τ_IE_* = 1. Also, *σ^EE^*, *σ^EI^*, *σ^II^* and the potential *V* are now functions of a single variable, *σ^IE^*. Therefore, the differential equation for *σ^IE^* can be written as

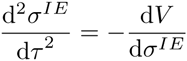

where

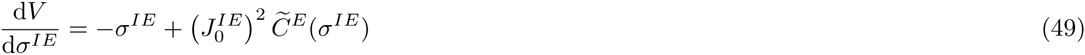

The instability of the fixed point occurs when, *V*′(*σ_IE_*) and *V*˝(*σ_IE_*), the first and the second derivatives of *V* with respect to *σ^IE^*, vanishes. Using Eq. (49) one has:

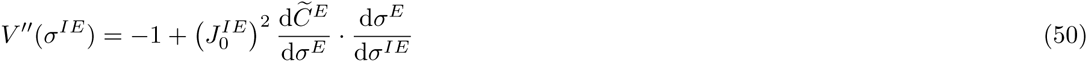

Since *σ ^α^* = *σ ^αE^* + *σ ^αI^*:

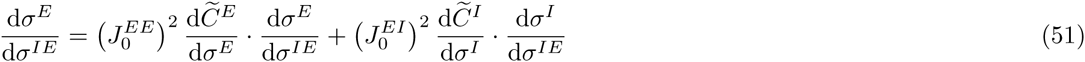

and

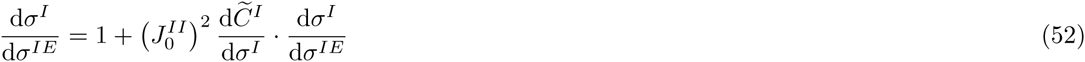

where

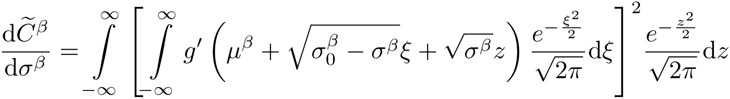

From Eqs. (51)-(52) one gets:

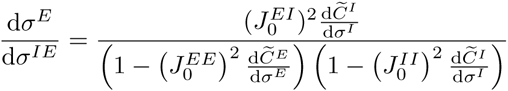

and:

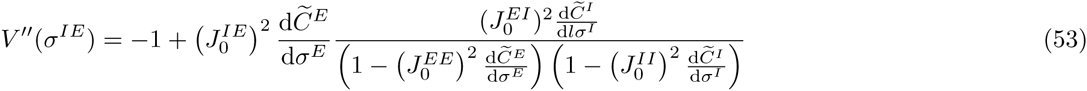

Thus at chaos onset, together with Eq. (41), 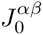, *σ ^α^* and *μ ^α^* obey:

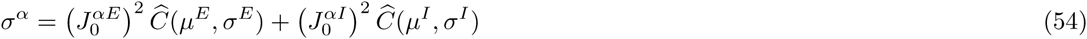

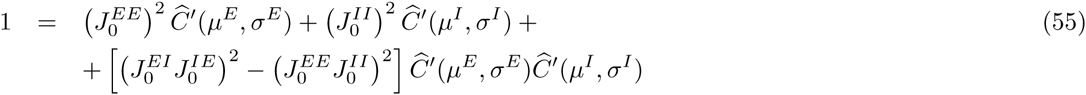

where:

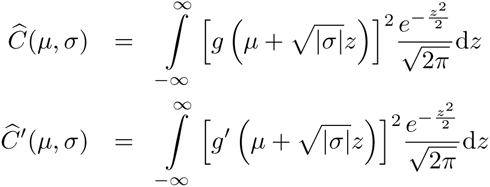

For instance for the threshold-linear transfer function we have

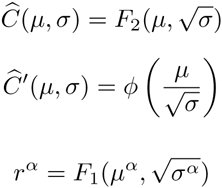

and where *F*_*i*_(*a, b*) are defined in equation (28).

It should be noted that if the transition to chaos occurs for the same parameters for which the fixed point loses stability and that this is controlled by a real eigenvalue crossing zero, the location of the transition will not depend on the synaptic time constant. If this is the case, Eq. (54) will characterize the location of the transition to chaos in the parameter space of the network in general and not only under the assumption of the separation of time scales under which we have established this condition.

#### On the stability of the fixed point

Let us denote the fixed point solution of the dynamics, Eq. (17), by: 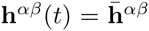. Writing 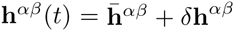 with 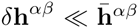, linearizing the dynamics and looking for solution of the form *δ***h** ∝ *e*^*λt*^) one gets:

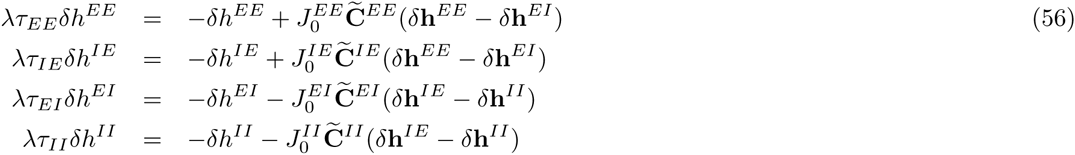

where the 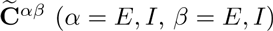 *N × N* sparse matrices with elements

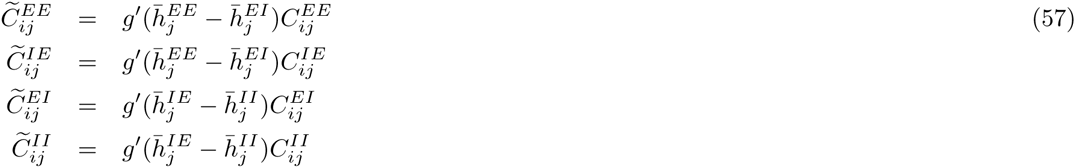

(**C** ^*α β*^ is the matrix of connectivity between populations *β* (presynaptic) and *α*). We are interested in instability onsets at which a real eigenvalue crosses 0.

Using Eqs. (56), it is straightforward to show that such an instability happens if the synaptic strength are such that:

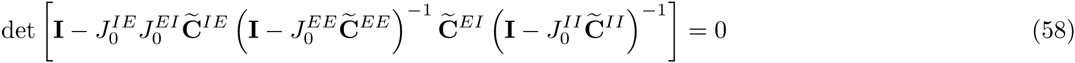

If 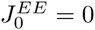, one can rewrite Eqs. (58) as:

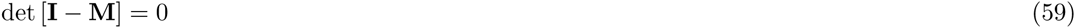

with:

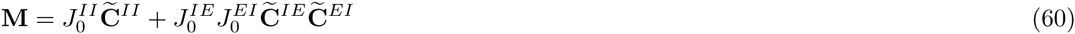

Let us assume that 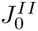 is fixed and such that for small enough 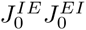 the fixed point is stable. When increasing, 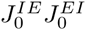 the fixed point loses stability when the value of 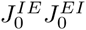 is the smallest for which Eq. (59) is satisfied, that is for which the largest real eigenvalue, *λ*_max_ of the matrix *M* crosses 1. If this instability also corresponds to chaos onset, Eq. (54), this would imply that the condition *λ*_max_ = 1 is equivalent to:

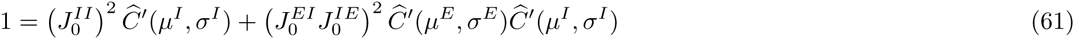

Interestingly, this condition means that the variance of the elements of the matrix 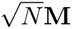 is equal to one leading us to conjecture that more generally the eigenvalue of the latter which has the largest real part and is given by:

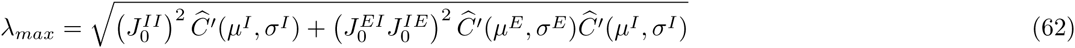

### Numerical simulations

#### Integration of network dynamics and mean-field equation solutions

The integration of differential equations, Eq. (15) and Eq. (18) (Eq. (3) in main text), was performed with a C code using the Euler method with fixed Δ*t* = *τ*_syn_/20 (the validity of the results was verified using smaller values of Δ*t*).

Simulations of the LIF spiking networks were done using a second-order Runge-Kutta integration scheme supplemented by interpolation of spike times as detailed in[66]. In all the spiking network simulations the time step was Δ*t* = 0.1 ms.

Self-consistent mean-field equations were solved with MATLAB function *fsolve*, which implements a ‘trust-region-dogleg’ algorithm or the Levenberg-Marquardt algorithm for non-square systems. Numerical calculations of integrals was done with MATLAB function *trapz*.

#### Population-averaged autocovariance

The population average autocovariance (PAC) functions of neuronal quantities *f*_*i*_(*t*) (*i* = 1*…N*) were computed as

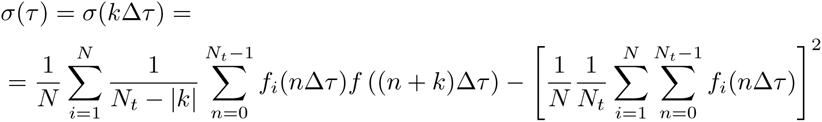

where *N*_*t*_ is the number of time samples for the calculation ofthe PAC. In all figures *f*_*i*_(*t*) = *h*_*i*_(*t*) except in Fig 16 where 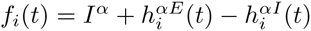. All PACs of spiking networks were calculated over 163.84 sec, and averaged over 10 realizations of the connectivity. For models Eq. (15) and Eq. (18), PACs were calculated over 2048*τ_syn_* after discarding 200*τ_syn_* of transient dynamics and averaged over 8 realizations.

#### Largest Lyapunov exponents

To calculate the maximal Lyapunov exponent, Λ, of the inhibitory network, Eq. (3), we simulated the system for a sufficiently long duration (200*τ_syn_*) so that it settled on the attractor of the dynamics. Denoting by 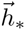 the network state at that time, we then ran two copies of the dynamics, one with initial conditions 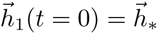 and the other with slightly perturbed initial conditions, 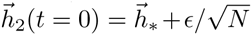 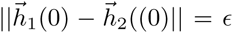, where ∥ ⋅ ∥ is the *l* ^2^ norm).Monitoring the difference, 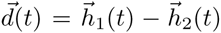 we computed 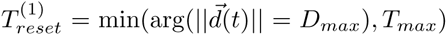 and 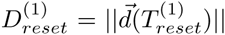 We then reinitialized the dynamics of the second network copy to 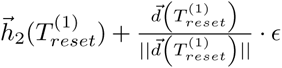 We iterated the process *n*times and estimate the Lyapunov exponent according to:

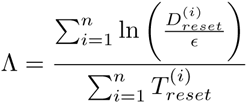

A similar method was used for two population networks, Eq. (15), the only difference being that the vector 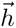 now had dimension 4*N*. Throughout the article we take *n*= 100, T_*max*_ = 5*τ*_*syn*_, *D*_*max*_ = 10^−3^ and *ϵ* = 10^−6^. The Lyapunov exponent values reported in this article are averages over 5 realizations of the networks.

#### Fraction of networks with a stable fixed point in rate dynamics

Fig. 10D in the main text plots the lines in the *J*_0_ − *I*_0_ phase diagrams of the threshold-power law rate model, for which 5%, 50%, 95% of randomly chosen networks have dynamics which converge to a fixed point. To compute these lines we simulated, for each value of *γ* and *J*_0_, 100 realizations of the network. For each realization, we computed the population average of the temporal variance the synaptic inputs, *ρ*:

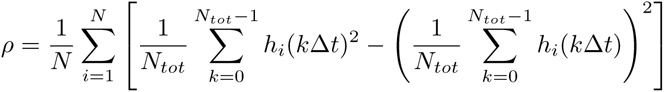

where *N*_*tot*_ is the total number of time steps of the simulations after discarding a transient with a duration of 256*τ_syn_*. The fixed point was considered to be unstable if *ρ >* 10 ^-9^. The fraction of unstable networks, *F*^*u*^, was fitted with a logistic function: *F*_*u*_(*J*_0_) = 100 [1 + exp (-(*J*_0_ *-J_m_*)/Δ *J*)]^−1^. The thick red line and red dots plot the values of *J*_*m*_ vs. *γ*, and the dashed lines are the values of *J*_0_ for which *F*_*u*_ = 95 and *F*_*u*_ = 5.

## Acknowledgments

We thank Gerard Benarous, Ran Darshan, Gianluigi Mongillo, Carl van Vreeswijk and Fred Wolf for insightful discussions. We thank Andrea Crisanti for sharing with us unpublished notes related to [19].

**Supporting Information Legends**

**S1 Text.** Finite size effects in inhibitory rate networks with sigmoid or threshold-linear transfer functions.

**S2 Text.** Chaos onset in inhibitory rate models with twice differentiable transfer functions.

**S3 Text.** Asynchronous rate chaos in non-leaky integrate-and-fire networks.

**S4 Text.** Onset of chaos in inhibitory rate models with threshold-power-law transfer functions.

**S5 Text.** Maximum Lyapunov exponents in the inhibitory LIF rate model.

**S6 Text.** Firing statistics in the inhibitory LIF spiking network.

**S7 Text.** Two-population rate model with a threshold-linear transfer function with 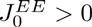.

**S8 Text.** The two mechanisms underlying asynchronous chaos in the two-population rate model with a threshold-linear transfer function.

**S9 Text.** The two mechanisms underlying asynchronous chaos in two-population LIF networks: Results for inhibitory neurons.

**S10 Text.** Two-population LIF rate and spiking models: In the II mechanism the PAC depends very mildly on *τ_IE_*.

**S11 Text.** Two-population LIF rate and spiking models: In the EIE mechanism the slow component of the PAC depends very mildly on *τ_II_*.

**S12 Text.** Two-population integrate-and-fire network with recurrent EE excitation, AMPA and NMDA synapses and fast inhibition.

**S1 Fig.** Finite size effects in inhibitory rate models with sigmoid or threshold-linear transfer functions.

**S2 Fig.** The potential function in inhibitory rate models with *g*(*x*) = *ϕ* (*x*).

**S3 Fig.** PACs in non-leaky integrate-and-fire networks.

**S4 Fig.** The potential function in inhibitory rate models with a threshold-linear transfer function.

**S5 Fig.** Lyapunov exponent in simulations of the inhibitory LIF rate model.

**S6 Fig.** Firing statistics in the LIF inhibitory spiking model.

**S7 Fig.** Two-population rate model with a threshold-linear transfer function and EE connections.

**S8 Fig.** The two mechanisms underlying chaos in the two-population rate model with a threshold-linear transfer functions.

**S9 Fig.** The PAC for the inhibitory neurons in the mechanisms underlying chaos in two-population LIF spiking and rate networks.

**S10 Fig.** The PAC depends very mildly on *τ_IE_* in the II mechanism.

**S11 Fig.** In the EIE mechanism the slow component of the PAC depends very mildly on *τ_II_*

**S12 Fig.** Two-population integrate-and-fire network with AMPA and NMDA synapses.

